# Crosstalk Between Phosphorylation and Ubiquitination Controls the Fate and Function of Strigolactone Signal Transducers

**DOI:** 10.1101/2025.08.27.672522

**Authors:** Lavanya Mittal, Neetu Verma, Dhanraj Singh, Shubhangi Pandey, Alok Krishna Sinha

## Abstract

Strigolactones (SL) are pivotal plant hormones that sculpt plant architecture by modulating shoot branching, root development, and meristem activity. While transcriptional responses downstream of SL perception have been well explored, the role of post-translational regulation fine-tuning these responses remains less understood. In this study, we identify a dual-layered regulatory module involving MPK4-mediated phosphorylation and MAX2-dependent ubiquitination that synergistically control the stability and function of BRC1, a key SL-responsive transcription factor. Phosphorylation by MPK4 stabilizes BRC1, enhancing its activity and SL sensitivity, whereas loss of phosphorylation leads to BRC1 degradation and functional inactivation. BRC1, in turn, directly activates *MPK4* transcription, establishing a positive feedback loop that amplifies SL signaling. Genetic analyses of the *brc1-2 × mpk4* double mutant reveals phenotypic defects and SL insensitivity additive to those observed in the *brc1-2* and *mpk4* single mutants, indicating that MPK4 and BRC1 act in parallel yet converging pathways downstream of SL. Additionally, MAX2 functions as a regulatory checkpoint that degrades non-phosphorylated MPK4 and BRC1, thereby resetting the signaling circuit to ensure accurate and timely response. Together, these findings illuminate a finely-tuned regulatory module integrating phosphorylation and ubiquitination to control the intensity and duration of SL responses, suggesting a model for hormone-driven developmental plasticity in plants.

## Introduction

For decades, strigolactones (SLs) were perceived primarily as rhizospheric signals, best known for inducing the germination of root-parasitic plants such as *Striga* and *Orobanche* (Cook et al., 1966). However, the perception of SLs underwent a major shift with the discovery of their endogenous roles in regulating plant development, marking their recognition as a new class of plant hormones. Among their best-characterised roles is the inhibition of axillary bud outgrowth, thus contributing to shoot branching control and overall plant architecture (Gomez-Roldan et al., 2008; Umehara et al., 2008; Ku et al., 2024). Since then, SLs have been shown to influence a broad range of developmental processes, including root architecture, leaf senescence, stem secondary growth, and environmental adaptation, particularly in response to phosphate deficiency and shade (Hu et al., 2020; Sun et al., 2022; Tian et al., 2022; Wang et al., 2022; Wu et al., 2022; Wang et al., 2023).

The molecular framework of SL biosynthesis and signaling has been extensively elucidated in recent years. They are derived from carotenoid precursors through a series of enzymatic steps involving DWARF27 (D27), two carotenoid cleavage dioxygenases, CCD7 and CCD8 and the cytochrome P450 enzyme, MORE AXILLARY GROWTH1 (MAX1), that convert carotenoid precursors into bioactive SLs (Alder et al., 2012; Al-Babili & Bouwmeester., 2015; Waters et al., 2017; Hu et al., 2020). SL perception occurs via a conserved signaling module comprising the α/β-hydrolase receptor DWARF14 (D14), the F-box protein MORE AXILLARY GROWTH2 (MAX2), and a group of transcriptional repressors belonging to the SMXL/D53 protein family (Wang et al., 2015; Yoneyama et al., 2018). Upon SL binding, D14 undergoes a conformational change that promotes its interaction with MAX2, leading to the formation of an SCF^MAX2^ E3 ubiquitin ligase complex that targets SMXL (SUPPRESSOR OF MAX-1 LIKE) proteins for proteasomal degradation (Soundappan et al., 2015; Mach, 2015; Shabek et al., 2018). This proteolysis releases transcriptional repression, allowing SL-responsive developmental and physiological programs to proceed. One such target is BRANCHED1 (BRC1), a TCP-family transcription factor that functions as a primary-response gene in the SL pathway. In axillary buds, SL-induced SMXL degradation de-represses BRC1 transcription within hours of treatment, indicating direct regulation (Braun et al., 2012; González-Grandío et al., 2017). Elevated BRC1 levels then suppress bud outgrowth, translating the early signaling event into inhibition of shoot branching (Aguilar-Martínez et al., 2007; Soundappan et al., 2015). Beyond shoot branching, BRC1 integrates additional hormonal and environmental cues. It promotes abscisic acid– mediated bud dormancy (González-Grandío et al., 2013), delays flowering under short-day conditions (Luo et al., 2019), and mediates cytokinin–auxin cross-talk to fine-tune meristem activity (Omoarelojie et al., 2019). It also influences leaf senescence by activating senescence-associated transcription factors under stress (Torres-Vera et al., 2014).

While this core mechanism functions effectively as a molecular switch that turns gene expression on or off, many developmental and environmental cues necessitate a more refined regulation, such as, modulating the strength, timing, or spatial distribution of gene activation to ensure precise control. Here, critical signal amplifiers and modulators such as Mitogen-Activated Protein Kinase (MAPK) cascades may serve as parallel signaling molecules that facilitate rapid and reversible responses to internal and external cues. Such modules may have the potential to relay SL cues from the membrane or cytoplasm directly to the nucleus, where they may phosphorylate transcription factors or other regulatory proteins, altering their stability, localisation, or activity (Meng & Zhang, 2013). Recent studies have begun to highlight the involvement of MAPK cascades as potential non-canonical routes that link SL perception to transcriptional responses (Jia et al., 2022; Ni et al., 2020), but such findings are still preliminary.

The MAPK cascade is a highly conserved signaling module that plays a central role in transducing a wide variety of extracellular and intracellular cues into appropriate cellular responses in plants. It operates through a three-tiered phosphorylation relay involving MAPKKKs, MAPKKs, and MAPKs, with activation requiring dual phosphorylation of conserved threonine and tyrosine residues (Sinha et al., 2011; Xu & Zhang, 2015; Bigeard et al., 2015; Jalmi et al., 2018; Manna et al., 2023; Singh et al., 2024). Activated MAPKs phosphorylate downstream targets, including transcription factors, enzymes, and cytoskeletal proteins, to control processes such as cell proliferation, differentiation, hormone signaling, and responses to biotic and abiotic stresses (Zhang and Zhang, 2022; Cristina et al., 2010). In *Arabidopsis thaliana*, the MAPK family consists of 20 members divided into four groups (A–D) based on the TXY activation motif (Ichimura et al., 2002). Groups A–C carry a TEY motif, while Group D features a TDY motif and divergent C-termini. MPK3 and MPK6 (Group A) are well studied for roles in immunity and development (Meng & Zhang, 2013), while MPK4 (Group B) is known for its functions in defense, cytokinesis and temperature responses (Petersen et al., 2000; Cristina et al., 2010; Verma et al., 2024). Other MAPKs, particularly in Groups C and D, remain largely uncharacterized (Bigeard et al., 2015).

Among the MPKs, MPK4 uniquely mirrors several SL-related functions. It is well known for roles in stress responses, cytokinesis, thermosensing, chromatin remodelling and hormone signaling (Bigeard et al., 2015; Verma et al., 2024). Notably, *mpk4* mutants display increased axillary bud outgrowth and impaired apical dominance - phenotypes resembling SL-deficient or insensitive lines such as *max2-1*, *smxl*, and *brc1-2* mutants (Gomez-Roldan et al., 2008; Soundappan et al., 2015). Furthermore, MPK4 has been shown to phosphorylate the transcription factor IPA1 (IDEAL PLANT ARCHITECTURE 1) under salt stress, stabilizing it and promoting its activity (Jia et al., 2022). IPA1 is a known downstream effector in SL signaling, playing a critical role in shoot architecture by modulating genes like *BRC1*.

Together, these observations led us to hypothesize that a parallel MAPK signaling pathway could potentially complement the canonical D14-SCF^MAX2^-SMXL signaling route, by adding an extra layer of regulation and flexibility in controlling downstream responses. This dual mode of signaling (targeted protein degradation coupled with phosphorylation-based modulation) would make SL signaling more robust and adaptable to multiple and fluctuating internal and external cues.

In this study, we demonstrate that MPK4 contributes to SL signaling by directly phosphorylating BRC1, serving as an additional regulatory layer to the canonical route. We further show that both MPK4 and BRC1 undergo ubiquitin-mediated degradation, revealing a coordinated system of post-translational control. Together, our results uncover a feedback-enabled mechanism in which phosphorylation and ubiquitination dynamically modulate SL signaling, expanding the current model of SL action in plant development.

## RESULTS

### MPK4 is upregulated and activated upon SL treatment

To investigate the role of the MAPK signaling cascade in linking SL perception to downstream gene expression, we focused on the expression dynamics of MPKs. Given their central role as signaling hubs, we systematically analyzed transcript levels of *MAPKs* representing three phylogenetic groups: Group A (MPK3, MPK6, MPK10), Group B (MPK4, MPK5, MPK11, MPK12, MPK13), and Group C (MPK1, MPK2, MPK7, MPK14). 7-day-old *Arabidopsis thaliana* (Col-0) seedlings were treated with 5 µM GR24 (synthetic SL) or DMSO (mock) over a time course (0, 1, 3, 6, 12 h). qRT-PCR revealed that six MAPKs— MPK1, MPK3, MPK4, MPK6, MPK13, and MPK14—responded to GR24 (Figure 1A and S1A). A change in expression upto 2 fold was considered as physiologically significant. To further delineate responsive MAPKs, we assessed their activation status through western blot analysis. Crude protein extracts from GR24- and mock-treated Col-0 seedlings were analysed using a phospho-specific pTEpY antibody, which detects the dually phosphorylated TEY motif - a marker of active MAPKs (Verma et al., 2021). A clear activation of ∼42 kDa MAPKs was observed at 6 h and 12 h post-GR24 treatment, in contrast to mock (Figure 1B).

**Figure 1.**
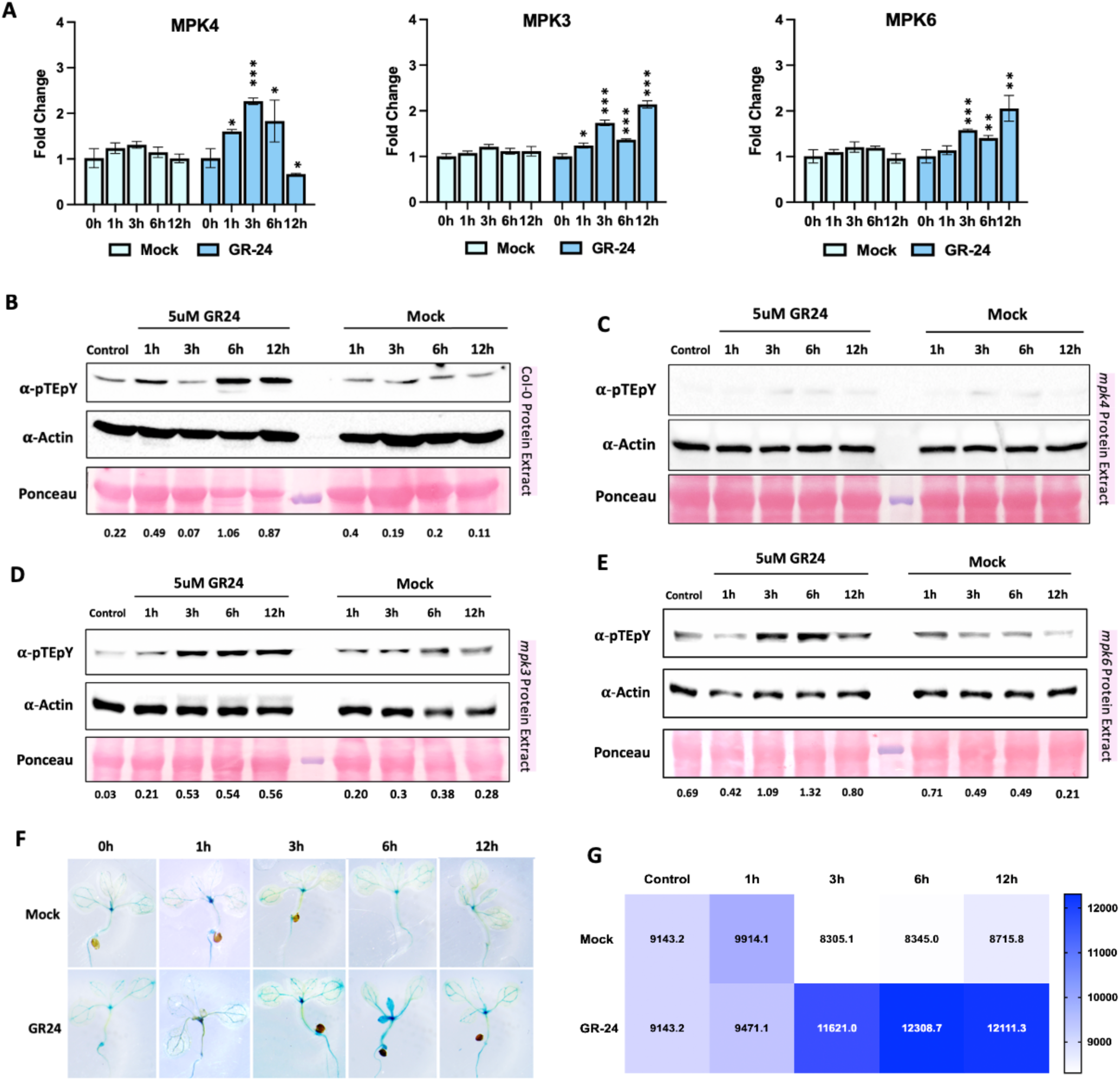
MPK4 is upregulated and activated upon SL treatment. **A.** Expression of group A, B and C MPKs in GR-24 and mock treated 7day old Col-0 seedlings. Values are means ± s.d. (n= 3). Expression of genes at each time point was compared to 0 h by Student’s t test; *p* value < 0.05 (*), < 0.005 (**), <0.0005 (***). **B, C, D and E.** Western blots from crude protein extracts of GR-24 and mock treated 7day old Col- 0 seedlings (B), *mpk4* mutant seedlings (C), *mpk3* mutant seedlings (D), and *mpk6* mutant seedlings (E), developed using anti-pTEpY antibody. Ponceau stainied blot shows RuBisCO protein. Actin protein was used as endogenous control. Numbers indicate the intensity of protein bands relative to Actin at each time point. **F.** Histochemical GUS staining of 7day old *proMPK4:GUS* seedlings grown in short day (SD) condition at 22°C treated with GR-24/mock at different time points **G.** Quantitative measurement of GUS activity in 7 day old *proMPK4:GUS* seedlings treated with GR-24. Activity is expressed in pmol of 4- MU/min/mg protein Values are means ± s.d. (n= 3).

Of the six GR24-responsive MAPKs - MPK3, MPK4, and MPK6 are among the most extensively characterized, known for roles in hormonal signaling, stress responses, and cell cycle regulation (Meng and Zhang 2013; Sethi et al., 2014; Verma et al., 2020). To identify the active kinases, phospho-blots were repeated in *mpk3, mpk4*, and *mpk6* mutants. Only *mpk4* lacked MAPK activation, suggesting MPK4 is essential for SL-induced MAPK activation (Figure 1C–E). These data suggest that MPK4 is not only transcriptionally induced but also uniquely activated by SL.

To corroborate the transcriptional upregulation of *MPK4* in response to SL, we monitored GUS activity in *proMPK4:GUS*seedlings treated with GR24 or mock as before. Increased GUS staining and activity were observed from 1 to 6 h post-GR24, declining slightly by 12 h, while mock controls showed only basal activity (Figure 1F, G). Western blot using MPK4-specific antibodies confirmed increased protein accumulation, peaking at 6 h after GR24 treatment (Figure S1B). Together, these findings demonstrate that MPK4 is both transcriptionally and post-translationally regulated in response to SL signaling. The coordinated increase in *MPK4* transcript and protein, coupled with its specific phosphorylation and the loss of activation in the *mpk4* mutant, propose MPK4 as a central, non-redundant component of the SL-induced MAPK cascade in *Arabidopsis*.

### MPK4 interacts with the SL response transcription factor BRC1

As a serine/threonine kinase, MPK4 may phosphorylate downstream transcriptional regulators or signaling components involved in SL responses. Six candidate genes- *WOX4, SPL13, SMXL6, BRC1, WRKY41*, and *PAP1* – were selected based on their roles in SL signaling, and the presence of conserved MPK phosphorylation motifs (SP/TP) (Bigeard et al., 2015; Verma et al., 2021) (Figure S2A). All selected candidates had at least one predicted site, suggesting they could be MPK4 substrates. However, only BRC1 had conserved SP/TP motifs across homologues and paralogues (Figure 2A, B). *BRC1* contained three predicted MAPK sites: S^34^P, S^65^P, and T^279^P, among which S^34^P and S^65^P were conserved, implying that T^279^P may be less critical for MPK4-mediated phosphorylation (Figure 2A, B). Next, we examined *BRC1* expression to evaluate whether its transcriptional response aligns with that of *MPK4* under SL treatment.. *BRC1* transcript levels mirrored *MPK4*, peaking at 6h post-GR24 treatment, suggesting a potential regulatory link between MPK4 and BRC1 in SL signalling (Figure 2C).

**Figure 2.**
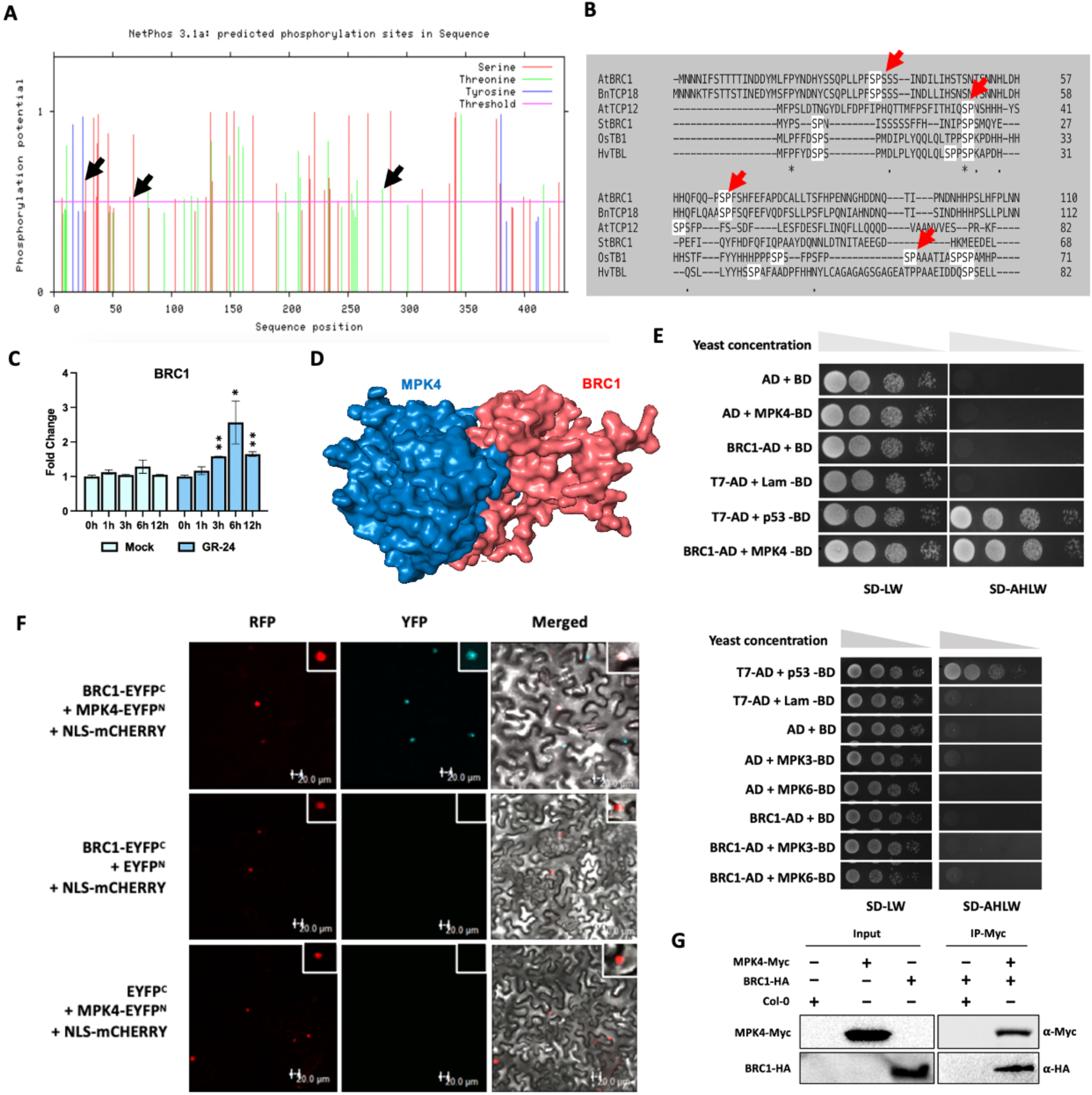
BRC1 interacts with MPK4. **A.** NetPhos3.1 results showing predicted MAPK phosphorylation motifs in BRC1 protein sequence. Black arrows indicate putative MAPK phosphorylation sites with score> 0.5 **B.** Multiple sequence alignment result of Clustal Omega using protein sequences of AtBRC1 and its homologues – BnTCP18 (*Brassica napus* TCP18), StBRC1 (*Solanum tuberosum* BRC1), OsTB1 (*Oryza sativa* TB1), HvTBL (*Hordeum vulgare* TB1-like) and the paralogue AtTCP12 (*Arabidopis thaliana* TCP12). Red arrows indicate conserved SP motifs. **C.** Transcript level of *BRC1* in GR-24 and mock treated, 7 day old Col-0 seedlings. Values are means ± s.d. (n= 3). Expression of genes at each time point was compared 0h by Student’s t test: *p* value < 0.05 (*), < 0.005 (**). **D.** *In-silico* docked structure of BRC1 (red) and MPK4 (blue) proteins. Docking was performed using Hdock and analyzed in PyMOL. **E.** Yeast-two-hybrid assay showing interaction of BRC1 with MPK4 (Upper), and with MPK3 and MPK6 (lower). SD-LW and SD-AHLW stand for supplement dropouts -leucine, -tryptophan and -adenine,-histidine,- leucine, -tryptophan, respectively. **F.** Confocal microscopy images of BiFC assay showing interaction of BRC1 with MPK4 performed transiently in *N.benthamiana* leaves. Inset on top right is the enlarged image of signal. Scale bar, 20 µm. **G.** Co-IP assay with protein crude extracts of 7 day old seedlings of transgenic Arabidopsis lines over-expressing *MPK4*-Myc and *BRC*1-HA. Blots were developed using anti-Myc (α-Myc) and anti-HA (α-HA) antibody.

To test if BRC1 is a direct MPK4 target, we first performed *in silico* protein–protein docking. The docking result predicted a stable interaction between BRC1 and MPK4 with a high confidence score (0.85), suggesting the possibility of direct binding (Figure S2 B–H and Figure 2D). Yeast two-hybrid (Y2H) assays confirmed a specific interaction of BRC1 with MPK4, not with MPK3 and MPK6 (Figure 2E). Bimolecular Fluorescence Complementation (BiFC) assays in *N. benthamiana* showed strong nuclear YFP signal from co-expression of MPK4-EYFP^N^ and BRC1-EYFP^C^, confirming specific interaction in planta (Figure 2F and S3). To further substantiate the interaction between MPK4 and BRC1 *in vivo*, we performed a co-immunoprecipitation (Co-IP) assay using Aarabidopsis transgenic lines overexpressing MPK4-Myc and an estradiol-inducible line expressing BRC1-HA. Estradiol induction was optimised according to González-Grandío et al. (2017) (Figure S3B). Total protein extracts from both lines were co-incubated, and immunoprecipitation of MPK4 was carried out using an anti-Myc antibody. The interaction of BRC1 with immunoprecipitated MPK4 was analyzed by immunoblotting with anti-HA antibody. A band at approximately 50 kDa, corresponding to BRC1-HA was detected (Figure 2G). No HA signal was detected in negative control indicating that BRC1 was not retained in absence of MPK4-Myc (Figure 2G). These results confirm. a specific *in vivo* interaction between MPK4 and BRC1. Altogether, *in silico*, *in vitro*, and *in vivo* assays confirm a specific MPK4–BRC1 interaction, identifying BRC1 as a likely MPK4 substrate in the SL signaling pathway.

### MPK4 Phosphorylates BRC1 at Ser34 and Ser65

Given MPK4’s kinase activity and its interaction with BRC1, we tested whether MPK4 directly phosphorylates BRC1. An *in vitro* kinase assay using bacterially-expressed MPK4-His and GST-BRC1 in the presence of [γ-³²P] ATP showed a clear radioactive signal at ∼75 kDa, confirming direct phosphorylation of BRC1 by MPK4 (Figure 3A). No phosphorylation was observed when MPK3 or MPK6 was used, indicating specificity to MPK4 (Figure S3C). All kinases phosphorylated MBP, validating their activity (Figure S3D). To identify the phosphorylation site(s), we conducted an *in vitro* kinase assay using MPK4-His and GST-BRC1, followed by immunoblotting with α-pSer and α-pThr antibodies. Only α-pSer antibody showed a clear signal at the expected size (∼75 kDa), whereas no detectable signal was observed with the α-pThr antibody, effectively ruling out T^279^ as a phosphorylation target (Figure 3B). We then mutated S^34^ and S^65^, individually and combinatorially to alanine due to its non-phosphorylatable nature using site directed mutagenesis (Figure S3E and Figure 3C). The mutated *BRC1* gene sequences confirmed by sequencing, and protein sequences are shown in Figure S4 A-H). *In vitro* kinase assays with these variants (WT, S^34^A, S^65^A, and S^34^A,S^65^A double mutant), revealed that substitution of S^34^ alone had a small impact on BRC1 phosphorylation, whereas mutation of S^65^ led to a marked reduction in the phosphorylation signal (Figure 3D). Strikingly, the double mutant (S^34^A,S^65^A) completely abolished phosphorylation, indicating that both S^34^ and S^65^ serve as phosphorylation targets of MPK4, with S^65^ playing a predominant role (Figure 3D).

**Figure 3.**
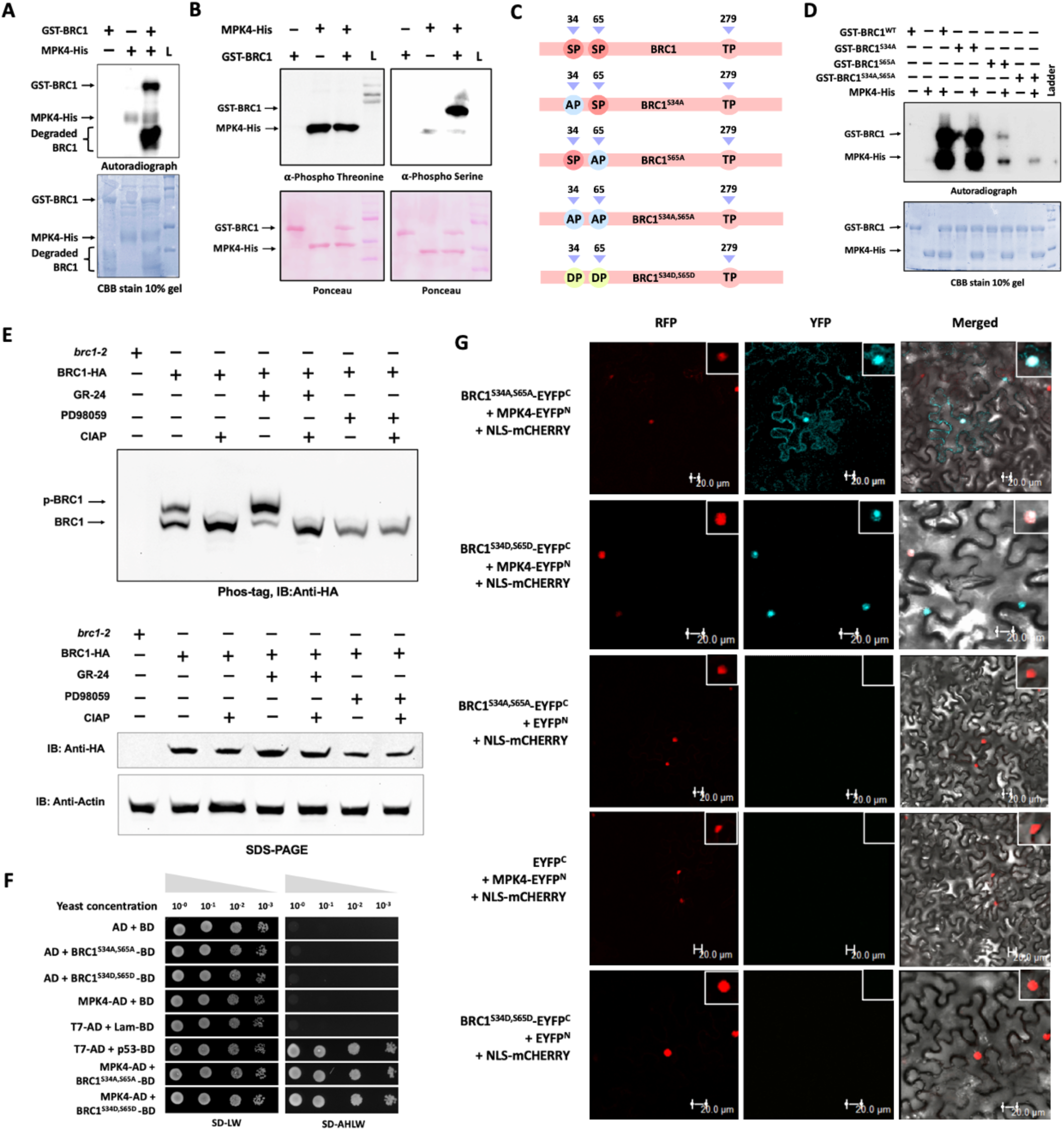
MPK4 phosphorylates BRC1 at Ser34 and Ser65. **A.** *In vitro* kinase assay showing phosphorylation of bacterially-expressed GST-BRC1 by MPK4-His. CBB (Commassie Brilliant Blue) stained 10% SDS page gel shows loaded proteins. **B.** Western blots of an *in vitro* kinase assay between bacterially-expressed GST-BRC1 and MPK4-His proteins. Ponceau stained western blot shows protein loading. Left blot is developed by anti-p-Threonine antibody and right blot is developed by anti-p-Serine antibody. Pictorial representation of all the *BRC1* mutant constructs designed through site-directed mutagenesis. **D.** *In vitro* kinase assay of various GST tagged mutant versions of BRC1 with MPK4-His. All proteins used are bacterially-expressed. CBB (Commassie Brilliant Blue) stained 10% SDS page gel shows loaded proteins. **E.** Western blots of *in-vivo* phosphorylation assay using 7% Phos-Tag gel (Upper panel) and 10% SDS-PAGE gel (lower panel). Blots were developed by anti-HA antibody. Actin was used as endogenous control. **F.** Yeast-two-hybrid assay showing interaction of MPK4 with BRC1^S34A,S65A^ and BRC1^S34D,S65D^. SD-LW and SD-AHLW stand for supplement dropouts -leucine, -tryptophan and -adenine,- histidine,- leucine, -tryptophan, respectively. **G.** Confocal microscopy images of BiFC assay showing interactions of MPK4 with BRC1^S34A,S65A^ and BRC1^S34D,S65D^ in *N.benthamiana* leaves. Inset on top right is the enlarged image of signal. Scale bar, 20 µm.

To confirm the phosphorylation of BRC1 by MPK4 *in vivo*, we used phos-tag mobility shift assay with BRC1-HA^Ind^ lines treated with GR24, PD98059 (MAPK inhibitor), or CIAP (phosphatase). In untreated BRC1-HA^Ind^ samples, two distinct bands were detected, corresponding to the phosphorylated and unphosphorylated forms of BRC1 (Figure 3E). Upon CIAP treatment, the band representing phosphorylated BRC1 was abolished, confirming its phosphorylation-dependent mobility shift. Notably, GR24 treatment led to an increased ratio of phosphorylated to unphosphorylated BRC1, indicative of enhanced phosphorylation upon SL signaling (Figure 3E). Under PD98059 treatment, only a single, faster-migrating band corresponding to the unphosphorylated form of BRC1 was detected, and its intensity was reduced compared to other treatments (Figure 3E).

Collectively, these results provide strong evidence that MPK4 phosphorylates BRC1 in response to SL signaling.

### Phosphorylation changes Localisation and Stability of BRC1

Having identified S^34^ and S^65^ as the phosphorylation targets of MPK4, we next sought to understand the functional consequence of this phosphorylation. To this end, we generated both phospho-dead BRC1^S34A,S65A^ and phospho-mimic BRC1^S34D,S65D^ versions of BRC1 (Figure S6 and S7). Aspartic acid was used to replace serine in phospho-mimic version due to its carboxyl group replicating the structural and electrostatic properties of a phosphorylated serine. These variants allowed us to dissect the physiological relevance of BRC1 phosphorylation by MPK4 in the context of SL signaling.

As an initial approach to investigate whether phosphorylation of the identified serine residues affects the interaction between BRC1 and MPK4, we performed a Y2H assay. Interestingly, MPK4 interacted with both modified forms of BRC1 (Figure 3F). BiFC assays in *N. benthamiana* confirmed this interaction *in vivo* (Figure 3G), suggesting that phosphorylation of BRC1 at S^34^ and S^65^ is not essential for its interaction with MPK4 and binding likely occurs independent of phosphorylation. However, the phospho-dead BRC1 localized with MPK4 in both nucleus and cytoplasm, while the phospho-mimic form remained nuclear, similar to the wild-type (Figure 3G and S5), suggesting that phosphorylation may play a role in restricting or refining the subcellular localization of the BRC1.

Phosphorylation by MAPKs often influences the stability and turnover of its substrates (Jagodzik et al., 2018; Ma and Nicolet, 2023). To explore whether phosphorylation affects the stability of BRC1, 7-day old estradiol inducible BRC1-GFP lines (BRC1-GFP^Ind^) were treated with PD98059 and DMSO (mock). Interestingly, in PD98059-treated seedlings, the GFP signal gradually increased until the 2-hour time point but then sharply declined compared to the mock (Figure 4A and 4B). One possible interpretation is that the early rise in GFP signal was driven by estradiol-induced expression, while the delayed effect of PD98059 allowed for initial protein accumulation. However, as the inhibitor progressively suppressed MAPK activity and thereby prevented BRC1 phosphorylation, the stability of unphosphorylated BRC1 may have decreased, leading to accelerated degradation. To further validate this observation, we performed western blot analysis using BRC1-HA^Ind^ lines treated similarly with PD98059, followed by harvesting at the same time points. Consistent with the confocal imaging data, we observed an initial increase in BRC1-HA protein levels, followed by a marked decrease after the 2-hour time point in the inhibitor-treated samples compared to the mock (Figure S6A). These observations suggest that MPK4-mediated phosphorylation could play a protective role in stabilizing BRC1 protein *in vivo*.

**Figure 4.**
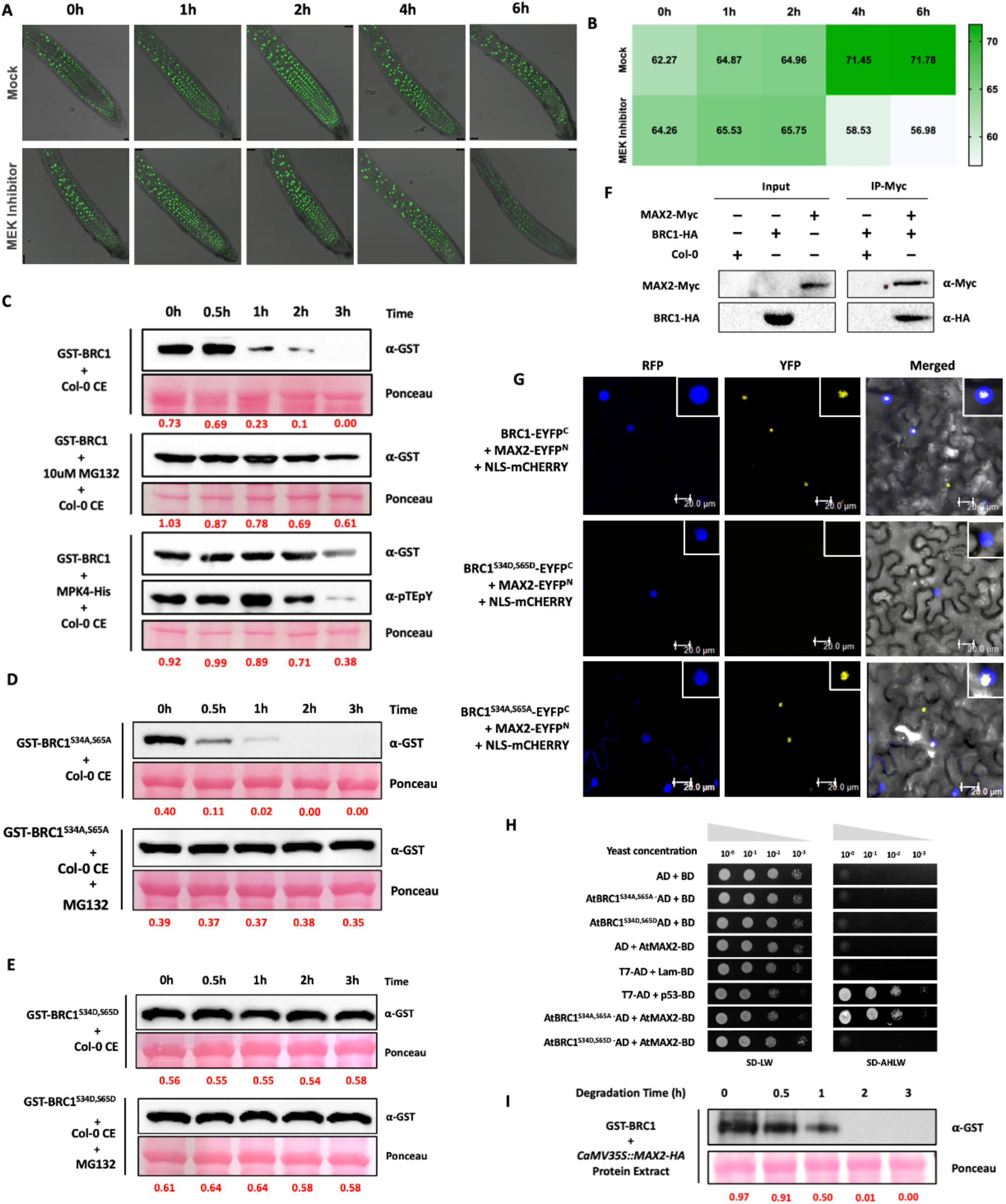
MAX2 interacts with and degrades unphosphorylated form of BRC1. **A.** Confocal microscopy images showing BRC1-GFP expression in root tips of *BRC1*-GFP^Ind^ lines treated with MEK inhibitor (PD98059) and mock. **B.** Pictorial representation of GFP signal intensity quantified in A. using the ImageJ software. **C.** Western blots showing cell-free degradation assays using 7 day old Col-0 protein crude extracts incubated with either GST-BRC1 alone (upper panel), with MG132 (middle panel) or with MPK4-His (Lower panel). **D.** Western blots showing cell-free degradation assays using 7 day old Col-0 protein crude extracts incubated with either GST-BRC1^S34D,S65D^ alone (upper panel) or with MG132 (lower panel). **E.** Western blots showing cell-free degradation assays using 7 day old Col-0 protein crude extracts incubated with either GST-BRC1^S34A,S65A^ alone (upper panel) or with MG132 (lower panel). (C, D and E) Western blots were developed using anti-GST antibody. Ponceau stained blot shows RuBisCo protein. Band intensities were quantified by ImageJ software. **F.** Co-Immunoprecipitation assay showing interaction between MAX2 and BRC1 using crude protein extracts of 7 day old seedlings of 35S:*MAX2*-Myc and estradiol inducible *BRC1*-HA^Ind^ lines. Western blots were developed by anti-Myc (α-Myc) and anti-HA (α-HA) antibodies. **G.** Confocal microscopy image showing BiFC assay of MAX2 with BRC1^WT^, BRC1^S34A,S65A,^ and BRC1^S34D,S65D^ transiently in *N. benthamiana* leaves. Inset on top right is the enlarged image of signal. Scale bar, 20 µm. **H.** Yeast-two-hybrid assay of MAX2 with BRC1^S34A,S65A^ and BRC1^S34D,S65D^ . SD-LW and SD-AHLW stand for supplement dropouts -leucine, -tryptophan and -adenine,- histidine,- leucine, -tryptophan, respectively. **I.** Western blot showing degradation of bacterially-expressed GST-BRC1 protein using crude protein extracts of *MAX2*-HA over-expression seedlings. Western blot was developed by anti-GST antibody. Ponceau stained blot shows RuBisCo protein. Band intensities were quantified by ImageJ software.

Since loss of protein stability is often due to targeted degradation via the ubiquitin–proteasome system (Lee et al., 2022; Langin et al., 2023), we performed cell-free degradation assays using bacterially-expressed GST-BRC1 incubated with Col-0 protein extracts. In one set of reactions, proteasome inhibitor MG132 was added. In parallel, a separate reaction included co-incubation with MPK4-His protein to facilitate *in vitro* phosphorylation of BRC1 prior to degradation. As expected, BRC1-GST alone underwent progressive degradation and was undetectable by 3 hours (Figure 4C). However, in the presence of MG132, BRC1 degradation was markedly reduced, indicating involvement of the 26S proteasome in its turnover. Pre-incubation with MPK4 also stabilized BRC1, suggesting phosphorylation protects it from degradation (Figure 4C). Next, we tested stability of phospho-dead (S^34^A,S^65^A) and phospho-mimic (S^34^D,S^65^D) BRC1 variants. The phospho-dead form was rapidly degraded within 1 hour, reversed by MG132 (Figure 4D). In contrast, the phospho-mimic variant remained stable regardless of MG132 treatment (Figure 4E).

Together, these findings reveal that phosphorylation of BRC1 at S^34^ and S^65^ by MPK4 influences its subcellular localization and enhances BRC1 protein stability by limiting its susceptibility to ubiquitin– proteasome-mediated degradation thus providing a mechanism for MPK4 to fine-tune BRC1 function in SL signaling.

### The F-Box Protein MAX2 Interacts with Unphosphorylated BRC1

MAX2, a key F-box protein in SL signaling, mediates ubiquitin-dependent degradation of SL-related proteins (Stirnberg et al., 2007; Jiang et al., 2013). Notably, many known substrates of MAX2 are either directly involved in SL-mediated developmental responses or serve as integrators of hormonal crosstalk with the SL pathway, such as SMXL proteins, D53-like repressors, and BES1 in brassinosteroid signaling (Wang et al., 2013; Wang et al., 2015). Given that unphosphorylated BRC1 is unstable, we hypothesized that MAX2 may target BRC1 for degradation.

To test this hypothesis, we examined whether BRC1 physically interacts with MAX2 - a prerequisite for ubiquitination. To this end we performed a Y2H assay and observed a direct interaction between BRC1 and MAX2 (Figure S6B). Co-IP using BRC1-HA^Ind^ and 35S::MAX2-Myc lines confirmed this interaction *in vivo* (Figure 4F). BiFC assays further validated their interaction in *N. benthamiana*, with fluorescence localized to the nucleolus (Figure 4G, S6C).

To further dissect the molecular mechanism linking phosphorylation and ubiquitination, we tested MAX2 binding to BRC1 phospho-variants. Unlike the phospho-dead version, phospho-mimic BRC1 failed to interact with MAX2 in both Y2H (Figure 4H) and BiFC assays(Figure 4G, S6C). Finally, degradation assays using bacterially-expressed GST-BRC1 incubated with extracts from *MAX2* overexpressing seedlings showed rapid degradation of BRC1 (Figure 4I), mimicking the phospho-dead variant’s instability as observed in Figure 4D.

Together, these findings provide compelling evidence for a dual-layered post-translational regulatory mechanism in which MPK4-mediated phosphorylation stabilizes BRC1 by preventing its ubiquitination and proteasomal degradation through the SCF^MAX2^ complex.

### MAX2 Interacts with Unphosphorylated MPK4 and Targets it for Degradation

MAX2, a central E3 ubiquitin ligase in SL signaling, targets many proteins for degradation. Given MPK4’s emerging role in this pathway, we hypothesized that MPK4 itself might be subject to regulation by MAX2. To explore this possibility, we first examined the potential interaction between MAX2 and MPK4 using an *in silico* approach. The predicted three-dimensional structure of MAX2 was obtained from the AlphaFold Database (Figure S7A and B). Different domains in MAX2 were predicted *in silico* (Figure S7C) and the positions of these domains have been shown in Figure 5A. *In silico* docking of predicted MAX2 structure with MPK4 revealed that MAX2’s F-box domain is positioned near MPK4’s TEY activation motif (<5 Å), a site of dual phosphorylation critical for MPK4 activation (Figure 5B). The proximity of the F-box domain to this critical regulatory site strongly suggests a functional interaction. Importantly, the addition of negatively charged phosphate groups at the TEY motif would likely introduce steric and electrostatic interference, potentially disrupting the interface with MAX2. This structural constraint supports a model in which phosphorylation of MPK4 not only activates its kinase function but may also prevent its recognition by the SCF^MAX2^ complex, thereby protecting it from ubiquitination and degradation. Further, Y2H assays confirmed physical interaction between MPK4 and MAX2 in yeast (Figure S8A). Co-IP using protein extracts from MAX2-HA and MPK4-Myc overexpression lines further validated this interaction (Figure 5C).

**Figure 5.**
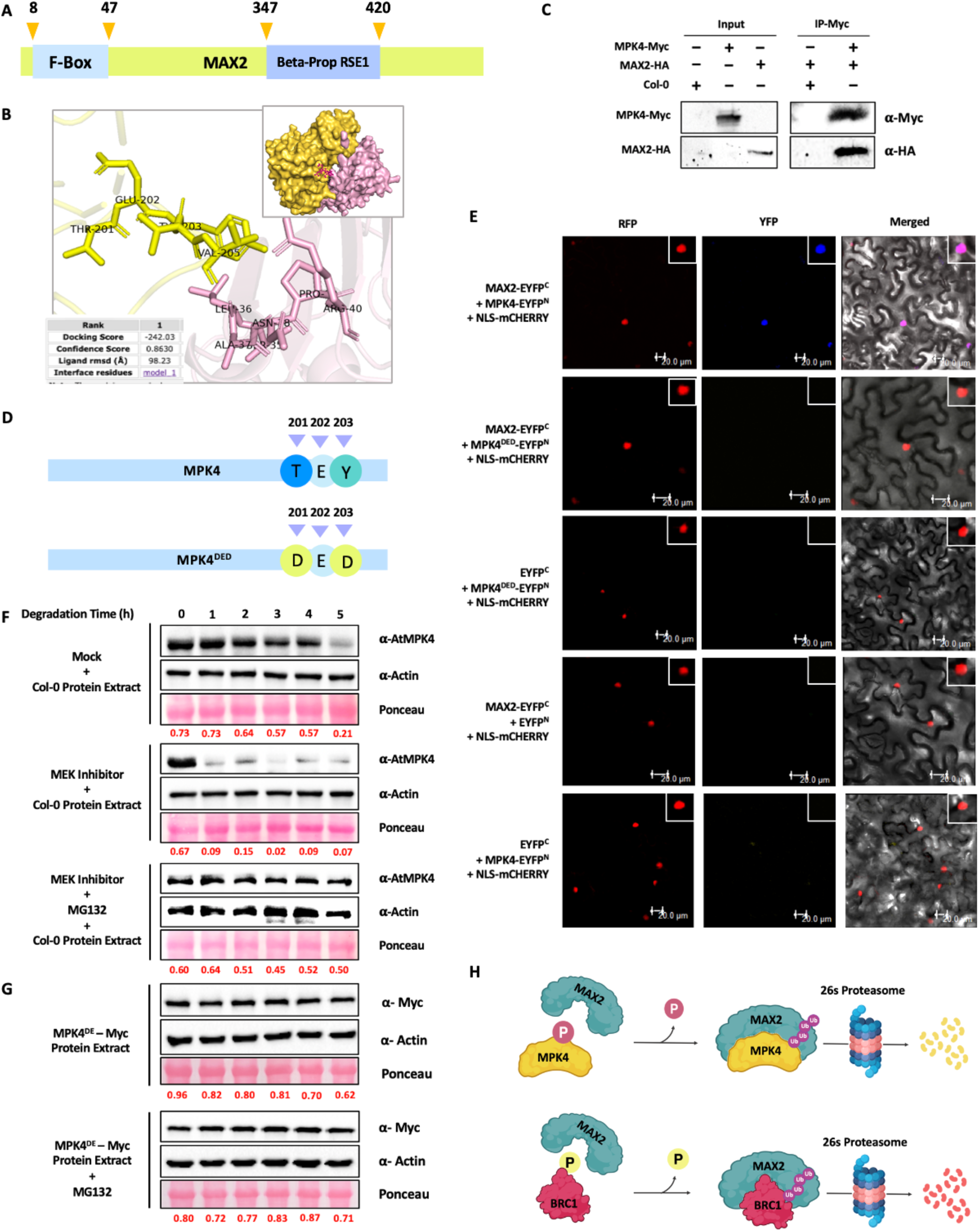
MAX2 interacts with and degrades unphosphorylated form of MPK4. **A.** Pictorial representation of predicted domains in MAX2 protein sequence. **B.** *In-silico* docking of predicted protein structures of MAX2 and MPK4 with an enlarged view of interacting residues of docked proteins of MAX2 and MPK4 (lower left panel). **C.** Co-Immunoprecipitation assay from protein crude extracts of 5 day old seedlings of *MAX2*-HA and *MPK4*-Myc over-expression lines. Western blots were developed by anti-Myc (α-myc) and anti-HA (α-HA) antibodies. **D.** Pictorial representation of amino acid residues mutated by site-directed mutagenesis in the protein sequence of MPK4. **E.** Confocal microscopy image of BiFC assay showing interaction of MAX2 with MPK4^WT^ and MPK4^DED^ transiently in *N.benthamiana* leaves. Inset on top right is the enlarged image of signal. Scale bar, 20 µm **F.** Western blots showing cell-free degradation of MPK4 protein using 7 day old Col-0 crude protein extracts treated with either DMSO (upper panel), with MEK inhibitor (middle panel) or with MG132 (Lower panel). Western blots were developed by anti-MPK4 antibody. Ponceau stained blot shows RuBisCo protein. Actin was used as loading control. Band intensities ratio of anti-MPK4 and anti-Actin is calculated using imageJ and depicted in red colour. **G.** Western blots showing cell-free degradation of MPK4^DE^ protein using crude protein extracts of 7 day old *MPK4*^DE^-Myc seedlings treated with either DMSO (upper panel), or with MG132 (Lower panel). Western blots were developed by anti-Myc antibody. Ponceau stained blot shows RuBisCo protein. Actin was used as loading control. Band intensities ratio of anti-MPK4 and anti-Actin is calculated using imageJ and depicted in red color. **H.** Pictorial representation of structural and molecular mechanism of MAX2 mediated degradation of BRC1 and MPK4.

To test if phosphorylation affects this binding, we generated a phospho-mimic MPK4^DED^ mutant in which T and Y in TEY motif were replaced with aspartic acid (Figure 5D). Y2H assays showed that MPK4^DED^ failed to interact with MAX2 (Figure S8B). BiFC in *N. benthamiana* confirmed that wild-type MPK4 interacts with MAX2 in the nucleus, but MPK4^DED^ does not (Figure 5E). Furthermore, western blot using Col-0, *max2-1* and *MAX2*-OE lines showed that MPK4 protein levels are slightly greater in *max2-1* mutant and drastically reduced in *MAX2*-OE line, implying that MAX2 is involved in MPK4 protein stability (Figure S8C). These data indicate that MPK4 interacts with MAX2 and phosphorylation of TEY motif of MPK4 disrupts this interaction, likely protecting MPK4 from ubiquitination.

To determine whether unphosphorylated MPK4 is inherently less stable, we utilized a constitutively active version of MPK4 (MPK4^DE^), carrying two point mutations D218G and E202A that mimic a persistently phosphorylated state (Berriri et al., 2012). Cell-free degradation assays using protein extracts from Col-0 seedlings treated with PD98059 showed rapid MPK4 degradation (∼85% loss within 1 h) (Figure 5F). However, the addition of MG132 prevented this degradation (Figure 5F). Conversely, extracts from plants expressing constitutively active MPK4^DE^-Myc showed that MPK4^DE^ remained stable over 5 h with or without MG132 (Figure 5G).

Together, these findings establish phosphorylation as a key determinant of MPK4 stability through a mechanism similar to BRC1: unphosphorylated MPK4 is targeted for SCF^MAX2^–mediated ubiquitin-proteasomal degradation, while phosphorylated or constitutively active MPK4 escapes degradation (Figure 5H). Thus, phosphorylation not only activates MPK4 but also shields it from proteasomal turnover, revealing a dual regulatory mechanism.

### Positive Feedback Regulation Between BRC1 and MPK4

Several reports showed that transcription factors that are phosphorylated by MAPKs can reciprocally bind to the promoters of their upstream kinases, modulating their expression either positively or negatively (Sethi et al.,2014; Singh and Sinha, 2016;). Therefore, we investigated whether BRC1 directly influences the SL-induced upregulation of MPK4, thereby establishing a self-reinforcing regulatory module within the SL signaling pathway. Firstly, we examined the *MPK4* promoter for the presence of BRC1 binding motifs. TCP transcription factors, including BRC1, lack a clear consensus DNA-binding motif. While some bind GGNCCC sequences, none were found in the *MPK4* promoter. We thus examined canonical bHLH motifs— E-boxes (CANNTG) and G-boxes (CACGTG)—as potential binding sites. The 1124 bp *MPK4* promoter was divided into three ∼300 bp fragments: *pMPK4-1, pMPK4-2,* and *pMPK4-3*. Two E-box motifs were located at −301 bp (*pMPK4-1*) and −705 bp (*pMPK4-2*) (Figure 6A). Docking simulations showed strongest BRC1 binding to *pMPK4-2* (score 0.9) (Figure 6B, S9A). Structural analysis revealed the binding of BRC1 downstream of the E-box in *pMPK4-2* (Figure 6B). To experimentally validate these predictions, yeast one-hybrid (Y1H) assays were performed using three *MPK4* promoter fragments. Y1H assay revealed specific binding of BRC1 with *pMPK4-2* (Figure 6C). This was further supported by electrophoretic mobility shift assay (EMSA) using bacterially-expressed GST-BRC1 and a γ-ATP radio-labelled probe containing the predicted binding site “TGGTGCA”. A concentration-dependent shift was observed, confirming direct binding of BRC1 to the predicted binding site on *MPK4* promoter (Figure 6D). Site-directed mutagenesis of predicted binding motif (TGGTGCA → TTGCACA, Probe M1P) abolished binding, while mutation of a nearby E-box (CAAATG → CGAATC, Probe M2E) did not (Figure 6E), confirming BRC1 specifically binds the TGGTGCA motif. To examine this regulatory interaction *in planta*, dual-luciferase and GUS reporter gene assays were performed in *N. benthamiana*. Co-infiltration of *CaMV35S::BRC1-GFP* with *pMPK4::LUC* or *pMPK4::GUS* reporter constructs significantly enhanced reporter activity, confirming transcriptional activation by BRC1 (Figure 6F and 6G). Phosphorylation often modulates TF DNA binding; however, dual-luciferase assays with BRC1^WT^, phospho-dead (S^34^A, S^65^A), and phospho-mimic (S^34^D, S^65^D) variants showed no difference in *MPK4* promoter activation, indicating phosphorylation at these sites does not affect BRC1’s DNA binding (Figure S9 B and C).

**Figure 6.**
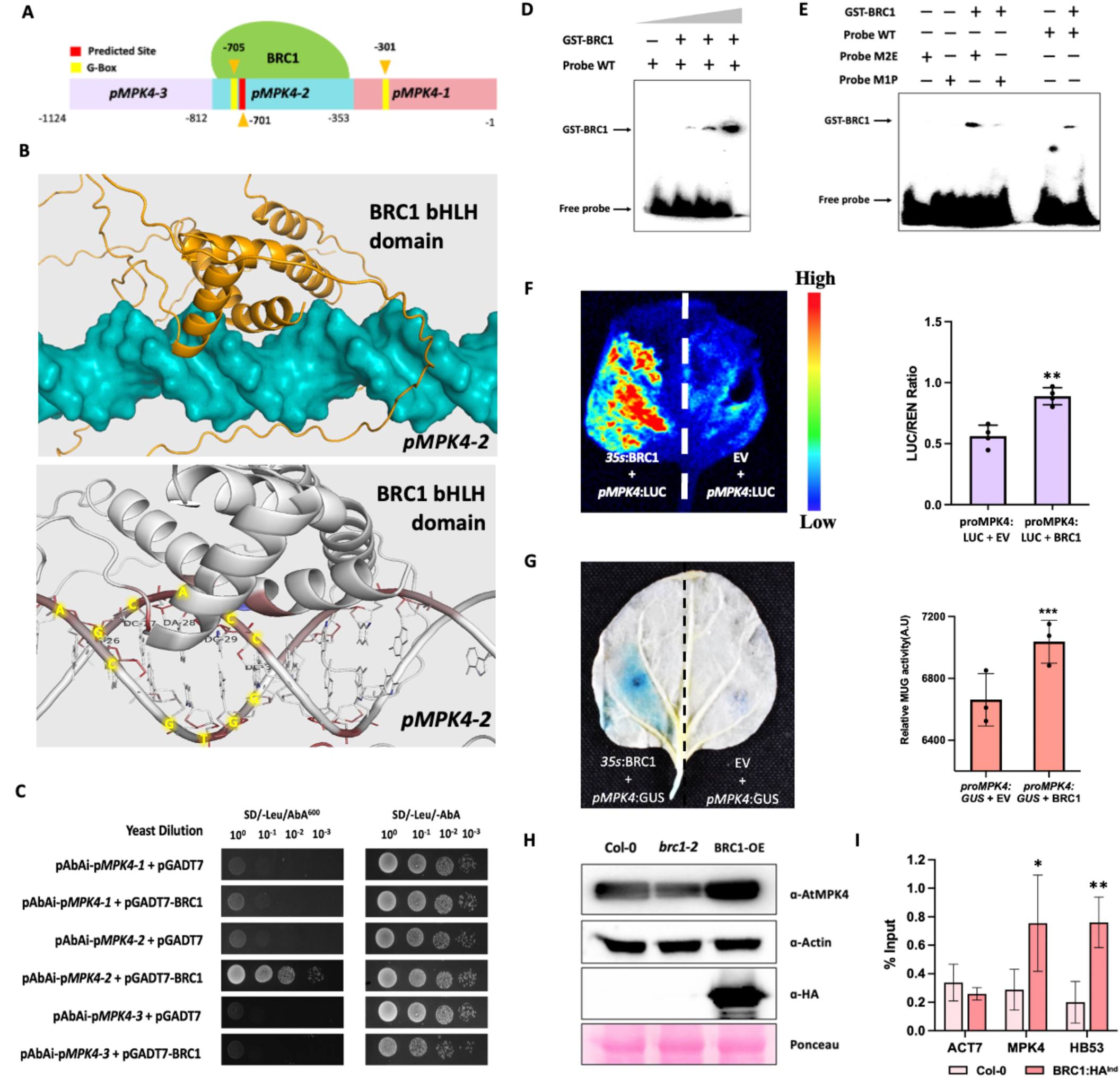
BRC1 binds to *MPK4* promoter and induces its expression. **A.** Pictorial representation of *MPK4* promoter depicting BRC1-binding site and promoter fragments used in various assays. **B.** *In-silico* docking of BRC1 on *pMPK4-2* fragment of *MPK4* promoter (upper panel). Docked structure of BRC1 on fragment *pMPK4-2* of *MPK4* promoter showing the nucleotide residues involved in binding (lower panel) **C.** Yeast-one-hybrid assay showing binding of BRC1 on *pMPK4-2* fragment. SD stands for supplement dropout. Leu and AbA stand for leucine and Aureobasidin A, respectively **D and E.** Electrophoretic mobility shift assays showing that BRC1 binds to the probe having predicted BRC1 binding site. M1P probe have mutated BRC1 binding site and M2E probe have mutation in a nearby E-box. **F.** Luciferace assay showing interaction between BRC1 and *pMPK4-2* fragment of *MPK4* promoter in transiently infiltrated *N. benthamiana* leaf (left panel). Quantification of Renilla luciferase activity in crude protein extracts of transiently infiltrated *N. benthamiana* leaves using luciferin as substrate (right panel). Values are means ± s.d. (n= 3). Significant difference was calculated by Student’s t test; *p* value < 0.0005 (***). **G.** GUS staining showing binding of BRC1 to *pMPK4-2* fragment in transiently infiltrated *N. benthamiana* leaf (left panel). Quantification of GUS activity in crude protein extracts of transiently infiltrated *N. benthamiana* leaves using MUG as substrate (right panel). Values are means ± s.d. (n= 3). Significant difference was calculated by Student’s t test; *p* value < 0.0005 (***) **H.** Western Blot using crude protein extracts of 7day old Col-0, *brc1-2* and *BRC1*-OE (*BRC1*-HA^Ind^) lines. Blots were developed by anti-MPK4 antibody. Actin was used as loading control. Blot developed with anti-HA antibody depicts protein levels of BRC1 in *BRC1*-HA inducible line. **I.** ChIP-qPCR to show the binding of BRC1 to the *MPK4* promoter in 7 day old Col-0 and *BRC1*-HA^Ind^ seedlings using anti-HA antibody. ChIP-qPCR of *HB53* and *Actin* promoters were used as positive and negative controls, respectively. Values are means ± s.d. (n= 6). Enrichment of each promoter fragment in *BRC1*-HA^Ind^ seedlings was compared to Col-0. Student’s t test; *p* value < 0.05 (*), < 0.005 (**).

Further, MPK4 transcript and protein levels were reduced in *brc1-2* mutants and elevated in estradiol-inducible *BRC1* overexpresson lines (Figure S9D, Figure 6H,), supporting positive regulation of *MPK4* by BRC1. ChIP-qPCR using anti-HA antibody in *BRC1*-HA plants demonstrated in planta binding of BRC1 to the *MPK4* promoter (Figure 6I).

Together, these data establish a positive feedback loop in which BRC1 directly binds to and activates *MPK4* promoter.

### BRC1 and MPK4 have Similar but Synergistic Effects on Plant Architecture

Histochemical GUS staining revealed a striking similarity in the tissue-specific expression patterns of BRC1 and MPK4 in promoter-driven GUS reporter lines (Figure S9E). Therefore, to understand the genetic relationship and physiological relevance of this regulatory loop, we analyzed the phenotypes of *brc1-2* and *mpk4* single mutants, and the *brc1-2 × mpk4* double mutant. Interestingly, analysis of various traits revealed overlapping roles of BRC1 and MPK4 in plant development. Both single mutants showed reduced height (∼27%) and increased branching (Figure 7A and 7B). The double mutant exhibited more severe, additive defects, including a ∼75% reduction in height and more increased branching (Figure 7A and 7B). Furthermore, SL sensitivity assays with 5 µM GR24 showed that both *brc1-2* and *mpk4* are partially insensitive, while the double mutant is largely unresponsive to SL. GR24 inhibited root elongation, lateral root development, and root hair formation in Col-0 but had diminished effects in single mutants and negligible effects in the double mutant (Figure 7 C-F). RAM length and meristematic cell number were also significantly higher in the double mutant (Figure 7G and 7H).

**Figure 7.**
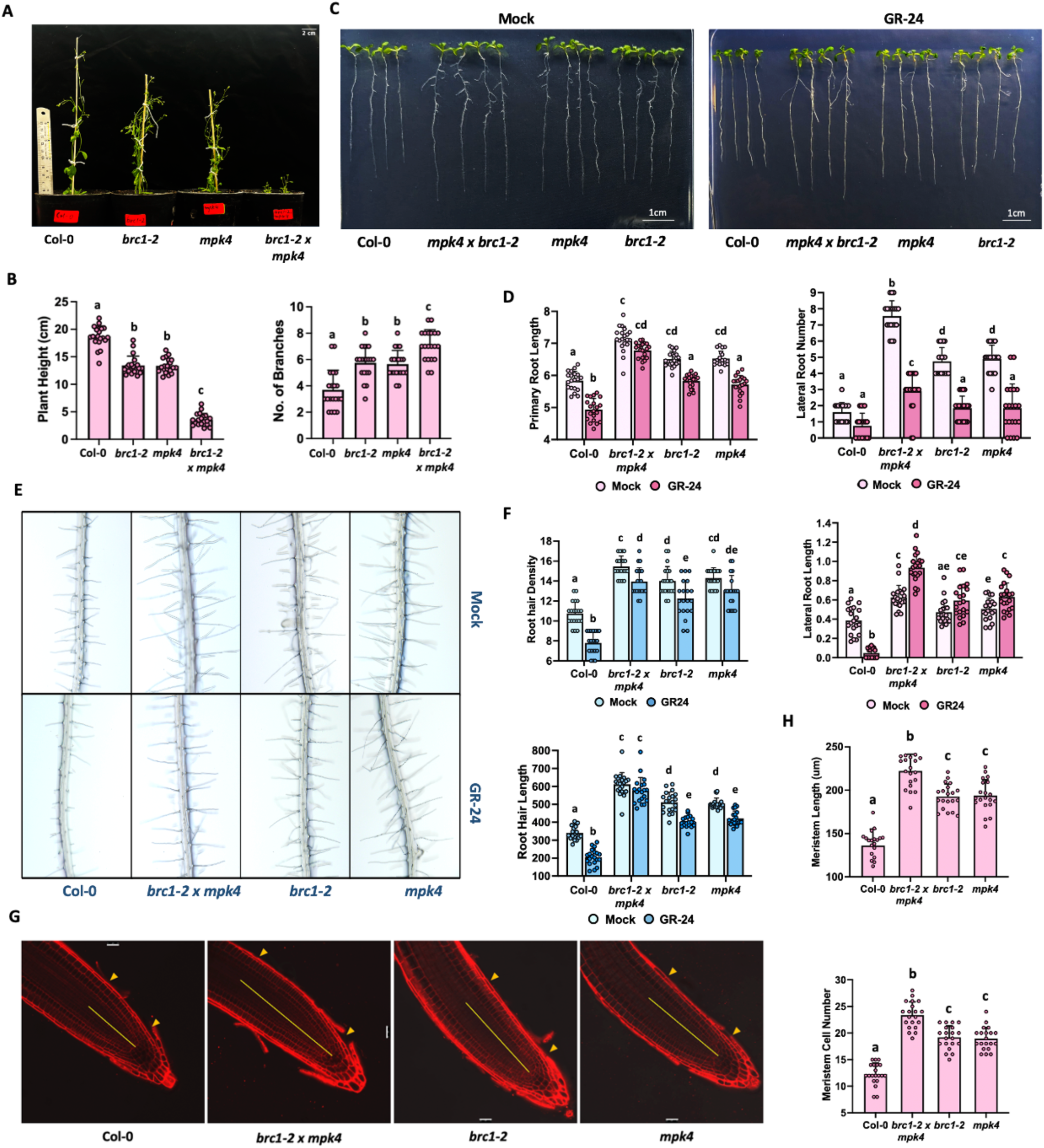
Functional relationship between BRC1 and MPK4. **A.** Image depicting the plant height and branching in 30 day old Arabidopsis plants grown in SD-condition. Bar = 2 cm (top right) **B.** Graph depicting plant height (cm) and number of primary branches in plants shown in A. Values are means ± s.d. (n= 20). Letters above the bars indicate significant differences (*p* < 0.05), calculated by One-way ANOVA with Tukey’s post hoc analysis. **C.** Images depicting the root phenotypes of 10 day old Arabidopsis seedlings given DMSO or 5uM GR-24 treatment for 5 days. Bar = 1 cm (bottom right). **D.** Graphs showing primary root length (left), lateral root number (right) and lateral root length (below right) of 10 day old Arabidopsis seedlings after mock/ GR-24 treatment. Values are means ± s.d. (n= 20). Letters above the bars indicate significant differences (*p* < 0.05), calculated by One-way ANOVA with Tukey’s post hoc analysis. **E.** Images showing root hairs in 10 day old Arabidopsis seedlings treated with DMSO or 5uM GR-25 for 5 days. **F.** Graphs showing root hair density and root hair length of 10 day old Arabidopsis seedlings treated with DMSO/5uM GR-24 (blue). Values are means ± s.d. (n= 20). Letters above the bars indicate significant differences (*p* < 0.05), calculated by One-way ANOVA with Tukey’s post hoc analysis. **G.** Confocal images showing 7 day old Arabidopsis roots stained with PI stain. Scale bar, 25µm. **H.** Graphs showing meristem length and meristem cell number in roots shown in G. Values are means ± s.d. (n= 20). Letters above the bars indicate significant differences (*p* < 0.05), calculated by One-way ANOVA with Tukey’s post hoc analysis.

These results indicate that BRC1 and MPK4 act additively to regulate plant architecture and SL responses. Their combined loss disrupts SL-mediated control of shoot and root development, highlighting their coordinated and potentially cooperative functions within the SL signaling pathway.

### Functional Relevance of MPK4 Mediated BRC1 Phosphorylation

To examine the physiological consequences of MPK4-mediated phosphorylation of BRC1, *brc1-2* plants were complemented with GFP-tagged BRC1^WT^, BRC1^S34A,S65A^ and BRC1^S34D,S65D^ each driven by the native BRC1 promoter (Figure S10A). Interestingly, the phosphodead lines resembled the *brc1-2* mutant, exhibiting reduced plant height, increased shoot branching, elongated primary roots, and an extended RAM (Figure 8 A-F). In contrast, the phosphomimic lines exhibited a phenotype reminiscent of BRC1 overexpression, including delayed bolting and flowering, indicative of a prolonged vegetative phase (Figure 8A and 8B). These lines also displayed a reduced RAM size (Figure 8C and 8D) and an enhanced sensitivity to GR24, consistent with increased SL responsiveness (Figure 8 E and F). Strikingly, upon GR24 treatment, BRC1^S34D,S65D^ lines showed a dramatic reduction in primary root length, whereas BRC1^S34A,S65A^ lines maintained longer primary roots than wild-type (Figure 8E and 8F). These observations suggest that phosphorylation status of BRC1 directly influences SL sensitivity and therefore plant architecture.

**Figure 8.**
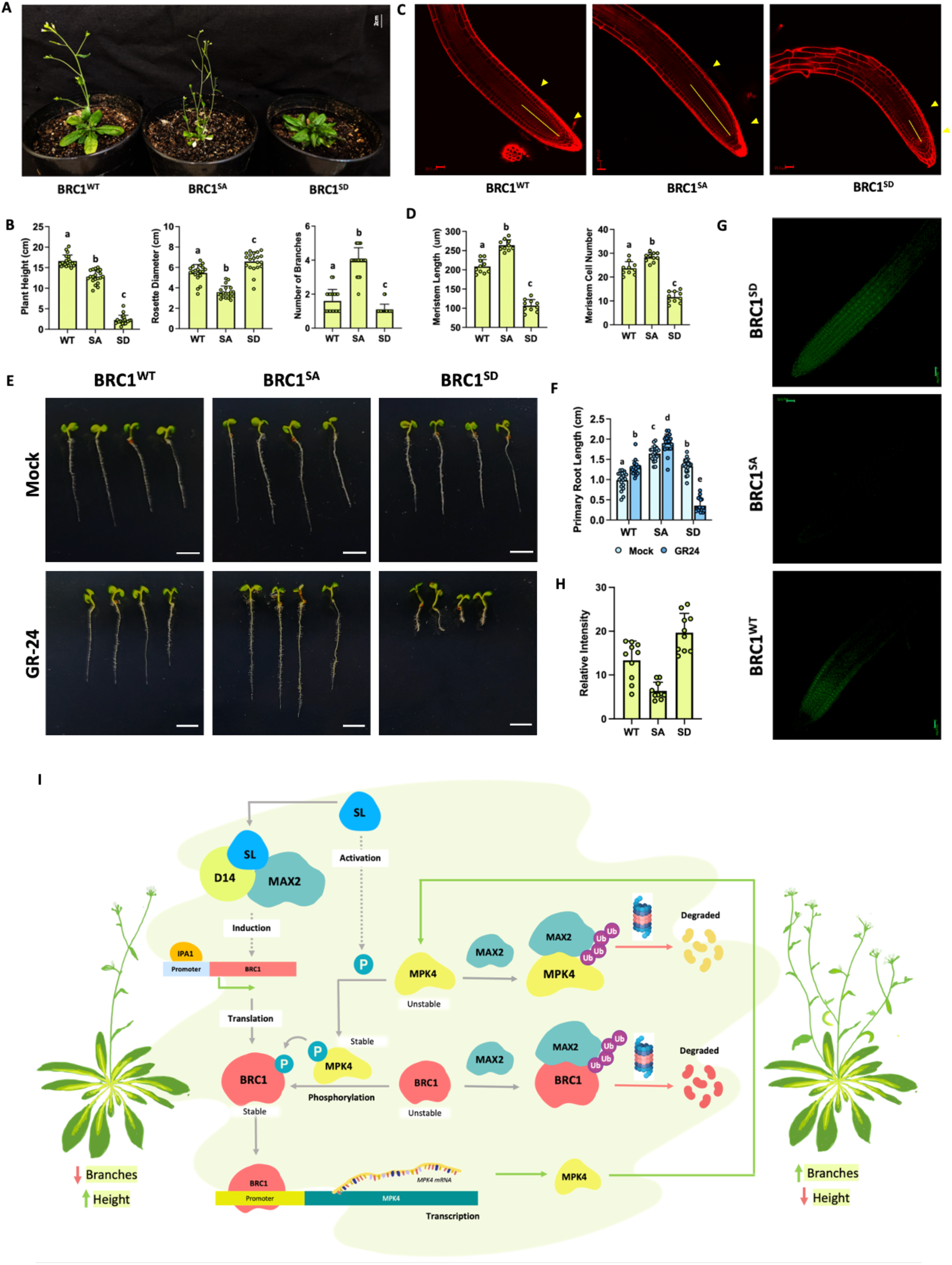
Phosphorylation of BRC1 is important for its function. **A.** Image depicting the plant height and branching in 25 day old Arabidopsis plants grown in SD-condition. Bar = 2 cm **B.** Graphs depicting plant height, rosette diameter, and number of primary branches in plants shown in A. Values are means ± s.d. (n= 20). Letters above the bars indicate significant differences (*p* < 0.05), calculated by One-way ANOVA with Tukey’s post hoc analysis. **C.** Confocal image showing 7 day old Arabidopsis roots stained with PI stain. Scale bar, 25um. **D.** Graphs showing meristem length and meristem cell number in roots. Values are means ± s.d. (n= 20). Letters above the bars indicate significant differences (*p* < 0.05), calculated by One-way ANOVA with Tukey’s post hoc analysis. **E.** Images depicting the root phenotypes of 7 day old Arabidopsis seedlings treated with DMSO or 5uM GR-24. Bar = 1 cm **F.** Graph showing primary root length after mock/GR-24 treatment. Values are means ± s.d. (n= 20). Letters above the bars indicate significant differences (*p* < 0.05), calculated by One-way ANOVA with Tukey’s post hoc analysis. **G.** Confocal images showing GFP signal in 7 day old Arabidopsis roots expressing GFP-tagged variants of BRC1. Scale bar, 25um. **H.** Graph showing relative GFP intensity in the roots shown in G. Values are means ± s.d. (n= 20). Letters above the bars indicate significant differences (*p* < 0.05), calculated by One-way ANOVA with Tukey’s post hoc analysis. **I.** Working model of regulation of SL mediated responses by dual-post-translational modification in Arabidopsis. Red lines indicate negative regulation. Green lines indicate positive regulation. Grey lines indicate neutral processes. Dotted lines indicate indirect processes.

To assess whether BRC1 protein dynamics are affected by phosphorylation, we visualized GFP-tagged BRC1 variants using confocal microscopy. Confocal imaging showed strong GFP signal in BRC1^WT^ roots, nearly absent signal in phospho-dead lines (suggesting protein destabilization), and enhanced but spatially restricted signal in phospho-mimic, likely reflecting increased stability within the smaller RAM (Figure 8G and 8H).

Taken together, these data reveal that phosphorylation by MPK4 is critical for the stability and function of BRC1 in planta. The phospho-mimic form behaves as a stabilized, possibly overactive variant of BRC1, resulting in enhanced SL sensitivity and suppressed RAM growth. In contrast, the phospho-dead version appears to be unstable and mimics a BRC1 loss-of-function phenotype.

## Discussion

Plants require precise and timely developmental responses to survive and thrive in ever-changing environmental conditions. To do so effectively, they rely on a well-coordinated arsenal of signaling molecules capable of rapidly transmitting information in response to both internal and external cues. Our study highlights a striking example of such a coordination by establishing MPK4 as a central component of the SL signaling pathway. We show that MPK4 functions as a key transducer of SL-specific hormonal signals working in close coordination with the D14-MAX2-SMXL signaling branch (Wang et al., 2020; Yoneyama et al., 2018). Specifically, MPK4 was found to be both transcriptionally induced and post-translationally activated following treatment with the SL analogue GR24 (Figure 1). Although our detailed time-course expression analysis revealed six members of MAPK family that displayed differential expression in response to SL treatment, MPK4 stood out due to its robust transcriptional upregulation coupled with strong activation. This conclusion is supported by the observation that MPK activation was abolished in the *mpk4* mutant but remained intact in *mpk3* and *mpk6* mutants, underscoring the non-redundant and essential role of MPK4 in SL signaling (Figure 1). This specificity is particularly noteworthy, given that MAPKs often function redundantly in stress responses (Wang et al., 2015; Singh et al., 2024). Although MPK3 and MPK6, both known players in hormone-related MAPK signaling, were transcriptionally induced by GR24, they were not activated at the post-translational level, which is a critical step required for transmitting the signal to downstream effectors (Bethke et al., 2009; Bigeard et al 2015). The preferential activation of MPK4 suggests the existence of a dedicated MAPK branch within the SL signaling network.

Activation of MPK4 upon SL treatment is expected to initiate critical phosphorylation events involving downstream regulatory proteins. This led to the next key question: which early SL-responsive transcription factor(s) does MPK4 target to propagate the signaling cascade? A combination of focused literature review and *in silico* analysis identified BRC1 as the most promising candidate. Notably, BRC1 harbours two high-confidence MAPK consensus motifs, S^34^P and S^65^P, both predicted with high confidence and highly conserved across its homologs and paralogs (Figure 2A and B). Functionally critical motifs such as kinase phosphorylation sites are typically preserved due to strong purifying selection, which acts to eliminate deleterious mutations that compromise essential developmental or signaling processes (Kimura, 1983; Lynch, 2007). The evolutionary persistence of these SP motifs across various plant lineages indicates they perform vital regulatory functions that have been maintained due to their importance. Moreover, BRC1 functions as a pivotal transcription factor that restricts the growth of axillary buds and plays a key role in shaping the plant’s branching pattern. It translates SL signals into gene expression changes that reduce meristem activity in side buds, making it a central component of the SL pathway (Aguilar-Martínez et al., 2007; Braun et al., 2012; Seale et al., 2017). Given its central regulatory role, BRC1 represents a biologically relevant and functionally significant target of MPK4 within the SL signaling cascade.

We also observed a striking overlap in the temporal expression profiles of MPK4 and BRC1 in response to GR24, suggesting a regulatory axis between these two proteins (Figure 2C). This kind of coordinated expression has also been reported in other hormone signaling systems, where kinases and their targets often show similar expression patterns (Singh et al., 2023; Singh et al., 2024). A combination of computational and experimental approaches provided strong evidence for a specific interaction between MPK4 and BRC1 (Figure 2). Importantly, BRC1 did not interact with other SL-responsive MAPKs such as MPK3 and MPK6, reinforcing the specificity of this kinase-substrate relationship. Moreover, this interaction was specifically localized in the nucleus, consistent with BRC1’s role as a transcription factor (Niwa et al., 2013). Our *in vitro* kinase assays further confirmed that MPK4 directly phosphorylates BRC1, with site-directed mutagenesis revealing that both S^34^ and S^65^ serve as critical phospho-acceptor residues (Figure 3). The complete loss of phosphorylation in the S^34^A,S^65^A mutant strongly indicates that these sites are jointly necessary for full phosphorylation, with S^65^ playing a dominant role. These findings are consistent with the modular nature of MAPK-substrate recognition, wherein multiple weakly conserved sites often contribute cumulatively to regulatory outcomes (Bigeard et al., 2015; Rayapuram et al., 2018). Importantly, Phos-tag gel assays provided strong *in vivo* evidence of BRC1 phosphorylation upon GR24 treatment. The phosphorylation-dependent band shift was abolished either by phosphatase treatment or by inhibition of MAPK signaling using the MEK inhibitor. These results not only confirm the role of MPK4 in mediating BRC1 phosphorylation but also demonstrate that this modification is induced in response to SL signaling. Such stimulus-dependent phosphorylation is a well-established mechanism for modulating transcription factor activity in hormone signal transduction pathways (Withers & Dong, 2017).

Phosphorylation of transcription factors such as BRC1 can modulate their functions through multiple non-mutually exclusive mechanisms, including alterations in protein stability, subcellular localization, DNA-binding affinity, and interactions with co-regulators (Hunter 2007, Wang et al., 2015; Zhang and zhang, 2022). Each of these effects may contribute to shape the transcriptional output in response to SL signaling. Given this mechanistic complexity, dissecting the specific molecular consequences of phosphorylation is essential to understand how upstream kinase activity is translated into defined developmental outcomes. While wild-type and phospho-mimic BRC1 localized predominantly to the nucleus, the phospho-dead version showed a striking re-localization to both nucleus and cytoplasm. This suggests that phosphorylation promotes nuclear retention of BRC1, a regulatory mechanism also observed in other MAPK substrates such as EIN3, WRKY33 and ABI5 (Yoo et al., 2008; Mao et al., 2011; Bhagat et al., 2025). Although the physical interaction between BRC1 and MPK4 was unaffected by phosphorylation status, reinforcing the idea that phosphorylation fine-tunes the functional compartmentalisation of BRC1 rather than its binding affinity for the kinase. Such spatial regulation may act as an additional control mechanism for restricting BRC1’s transcriptional activity to the nucleus under specific signaling conditions.

In addition to influencing localization, MPK4-mediated phosphorylation markedly enhanced BRC1 protein stability. Time-course experiments using inducible *BRC1*-GFP and *BRC1*-HA lines treated with the MEK inhibitor reveal that blocking MAPK activation leads to BRC1 destabilization followed by its degradation. The use of the proteasome inhibitor MG132 in our cell-free degradation assays, further confirmed that this degradation was dependent on the 26S proteasome, implicating ubiquitin-mediated turnover as the underlying mechanism (Figure 4). Phosphorylation thus appears to serve a protective function, shielding BRC1 from proteasomal degradation, a mechanism reminiscent of stabilization pathways seen in other hormone-related transcription factors such as BES1 and MYC2 (Zhai et al., 2013; Xing et al., 2023).

In many cases where phosphorylation regulates the stability of a protein whose degradation occurs via the 26S proteasome pathway, the potential involvement of F-box proteins often becomes a focal point. This is because F-box proteins, as substrate recognition subunits of SCF (Skp1-Cullin-F-box) E3 ubiquitin ligase complexes, typically recognize specific phosphorylated motifs on target proteins, known as phospho-degrons (Deshaies, 1999; Reed, 2005). Such phosphorylation-dependent recognition ensures precise temporal control over protein turnover in numerous signaling pathways. Therefore, it was reasonable to hypothesize that an F-box protein could mediate its ubiquitination. Among the known F-box proteins in Arabidopsis, MAX2 stands out as a critical regulator in strigolactone signaling, targeting a diverse set of transcriptional regulators for degradation. These include SMXL proteins (Nelson et al., 2011; Waters et al., 2012; Jiang et al., 2013), D53-like repressors (Wang et al., 2015; Soundappan et al., 2015), and BES1, a key component of brassinosteroid signaling (Wang et al., 2013). Additionally, MAX2 mediates the degradation of other transcriptional repressors involved in shoot branching regulation, underscoring its broad role in hormonal crosstalk and developmental control (Soundappan et al., 2015; Li et al., 2017). We identified a robust interaction between BRC1 and MAX2, especially localized to the nucleolus, providing initial molecular evidence supporting a direct regulatory relationship between the two proteins. We then connected this interaction to functional stability by showing that phospho-dead BRC1, which is prone to degradation, readily binds MAX2, whereas the phospho-mimic form fails to do so, suggesting that phosphorylation protects BRC1 from MAX2 recognition. Finally, our cell-free degradation assays using extracts from *MAX2*-overexpressing plants demonstrated accelerated degradation of BRC1, consistent with MAX2 mediating its turnover. Altogether, these results delineate a mechanism whereby MPK4-mediated phosphorylation stabilizes BRC1 by blocking its interaction with MAX2, thereby preventing ubiquitination and proteasomal degradation.

Building upon this framework, we extended our investigation to examine whether a similar mechanism governs the stability of MPK4 itself. Our interest in MPK4 was spurred by many reasons. First, MPK4 had already emerged as a crucial upstream kinase responsible for phosphorylating and stabilizing BRC1, thereby modulating a central transcriptional output of SL signaling. This positioned MPK4 as a regulatory node whose own stability could have significant downstream consequences. Second, our earlier results demonstrated that unphosphorylated BRC1 is selectively degraded via a MAX2-dependent pathway, prompting us to consider whether upstream elements like MPK4 might be similarly regulated to reinforce signaling fidelity. Third, many F-box proteins, including MAX2, are known to recognize phospho-degrons or unphosphorylated motifs in their targets, a mechanism that allows for dynamic, phosphorylation-sensitive degradation (Hunter, 2007; Skaar et al., 2013). Given that MPK4 activity is itself regulated by phosphorylation at the conserved TEY motif, it presented an ideal candidate for testing whether phosphorylation could also serve as a protective modification against proteasomal degradation. Moreover, prior reports have noted that MAPKs, including MPK4, can be subject to degradation under specific developmental or stress contexts (Lang et al., 2017), suggesting that their turnover might be regulated to facilitate the transient nature of hormonal cues. Our *in silico* docking analyses revealed that amino acid residues within the F-box domain of MAX2 lie in close spatial proximity, within ∼5 Å to the TEY motif of MPK4, a site essential for MPK4 activation via dual phosphorylation. This intimate alignment suggested that even subtle changes at the TEY site could disrupt the interaction. Given the high negative charge and bulky nature of phosphate groups, we reasoned that phosphorylation at the TEY motif could introduce steric hindrance and electrostatic repulsion, thereby obstructing MAX2’s access to MPK4. This premise was validated experimentally where the physical interaction between MPK4 and MAX2, was abolished when the TEY motif was mutated to phospho-mimic aspartic acid residues DED. These results presented a similar pattern to what we observed with BRC1, where phosphorylation conferred protection against MAX2-mediated degradation, suggesting a conserved mode of regulation within the SL signaling framework. Moreover, unphosphorylated MPK4 exhibited rapid degradation in a proteasome-dependent manner, especially upon inhibition of MAPK activation by PD98059. The stabilization of MPK4 in the presence of MG132 further confirmed the role of the 26S proteasome in this process. In contrast, the constitutively active MPK4^DE^ variant remained strikingly stable even in the absence of proteasome inhibition, highlighting that active or phosphorylated MPK4 evades MAX2-mediated turnover (Figure 5E). Together, these findings delineate a phosphorylation-dependent stability checkpoint for MPK4, whereby the active kinase state shields MPK4 from recognition and degradation by MAX2. This mirrors the regulatory mechanism observed for BRC1 and reveals a broader principle in which MAX2 acts as a surveillance factor that selectively targets inactive/unphosphorylated signaling components for degradation. Through this selective pruning, MAX2 ensures that only functionally competent effectors persist by swiftly eliminating inactive intermediates. Jia et al. (2022) demonstrated that MPK4 also phosphorylates IPA1, another key SL-responsive transcription factor. The study also reported that MPK4-mediated phosphorylation of IPA1 affects its protein stability. Since both BRC1 and MPK4 are targets of MAX2, it would be interesting to investigate whether IPA1 is also subject to MAX2-dependent degradation in a phosphorylation-sensitive manner. If true, it would advocate a broader regulatory principle at play, a model in which post-translational modifications, coordinated by kinases like MPK4 and E3 ligases like MAX2, may work in concert to fine-tune cellular responses to SL and potentially other hormonal cues. This regulatory precision may thus enable plants to initiate controlled hormonal responses while rapidly resetting the signaling machinery to preserve cellular homeostasis. In doing so, the system safeguards both the fidelity and the transient nature of hormonal signaling, allowing for dynamic yet controlled physiological outcomes. Moreover, this finding underscores the awe-inspiring collaboration of two important post-translational modifications - phosphorylation and ubiquitination, in orchestrating precise cellular signaling outcomes.

Given the critical role of MPK4 as a signaling kinase, understanding how its expression is transcriptionally regulated becomes equally important to grasp the complete regulatory circuit. Notably, several reports have demonstrated that MAPK substrates, particularly transcription factors phosphorylated by the kinase, can in turn regulate the transcription of the kinase itself, establishing transcriptional feedback loops (Mao et al., 2011; Birkenbihl et al., 2017). Such feedback mechanisms are essential for signal amplification, robustness, and fine-tuning of hormonal responses (Ichimura et al., 2000; Wang et al., 2022). Moreover, TCP transcription factors have been documented to participate in feedback regulation of MAPKs and other signaling kinases, binding directly to their promoters and modulate their expression (Martin-Trillo & Cubas, 2010; Viola et al., 2023; Meng & Zhang, 2013). Although BRC1 lacks a well-defined consensus binding motif and commonly reported TCP binding sites were absent in the *MPK4* promoter, we nonetheless focused on the two E-box elements present within the promoter sequence for *in silico* docking analysis. Interestingly, while BRC1 did not bind directly to the E-box motifs themselves, it showed a strong and specific interaction with a TGGTGCA site immediately downstream of one E-box within the *pMPK4-2* fragment. Moreover, *the brc1-2* mutant showed significantly reduced MPK4 transcript and protein levels, whereas overexpression lines exhibited enhanced *MPK4* expression, providing strong evidence of positive regulation in planta. Further confidence was gained through chromatin immunoprecipitation (ChIP) assays, where the enrichment of the *MPK4* promoter region by BRC1 binding was comparable to that of HB53, a well-established BRC1 target (Figure 6) (González-Grandío et al 2017). Collectively, these results demonstrate a positive feedback loop in which BRC1, acting downstream of SL signaling, directly binds to and activates *MPK4* expression, thereby reinforcing and amplifying the MPK4-mediated signaling branch.

Further, we used single mutants of *brc1-2* and *mpk4* and the corresponding double mutant (*brc1-2 × mpk4*) to elucidate the genetic relationship and functional hierarchy of this molecular association. Remarkably, both single mutants exhibited highly similar developmental defects, including reduced plant height, increased shoot branching, elongated primary roots, enhanced lateral root number, denser and longer root hairs, and an extended root apical meristem (RAM). These overlapping phenotypes suggest that MPK4 and BRC1 share common roles in modulating various facets of plant architecture and root system development. Strikingly, the double mutant exhibited significantly exacerbated phenotypes across all measured traits, pointing to an additive genetic interaction. This was particularly evident in GR24 sensitivity assays. In wild-type (*Col-0*) seedlings, GR24 treatment led to a pronounced suppression of primary and lateral root elongation, consistent with the established inhibitory role of strigolactones in root growth (Cheng et al., 2013; Sun et al., 2016). However, *brc1-2* and *mpk4* mutants exhibited partial resistance to GR24, and the *brc1-2 × mpk4* double mutant displayed near-complete insensitivity, suggesting a more profound disruption of SL signaling. While GR24 reduced root hair density and length in *Col-0*, these effects were only modestly mitigated in the single mutants and almost entirely lost in the double mutant. Notably, RAM length in the double mutant nearly doubled relative to *Col-0*, accompanied by a significant increase in meristematic cell number—again, with intermediate phenotypes in the single mutants (Figure 7). These observations are consistent with the role of BRC1 as a class II TCP transcription factor, known to limit meristematic activity by repressing genes involved in cell proliferation and growth (Martín-Trillo & Cubas, 2010; Nicolas & Cubas, 2016). The loss of BRC1 function leads to enhanced meristem size and cell division, and MPK4 appears to act in parallel or coordinately to enforce these developmental checkpoints. Collectively, these results support a model in which BRC1 and MPK4 function additively, yet non-redundantly, to regulate key aspects of plant growth, including shoot branching, root architecture, and RAM activity. Their combined loss significantly impairs the hormonal signaling that controls plant architecture, underscoring their critical, cooperative roles in the fine-tuning of SL-mediated developmental programs.

To elucidate how MPK4-mediated phosphorylation influences BRC1 function in plants, we investigated the developmental significance of this modification. While our previous results established that phosphorylation of BRC1 enhances its stability, these insights did not clarify the developmental consequences of this modification, particularly in the context of SL signaling. Phospho-dead and phospho-mimic complementation lines of BRC1 under its native promoter allowed us to functionally mimic absence or constitutive presence of a phosphate group at the identified sites. We could thus assess the direct impact of this modification on BRC1-driven developmental processes in Arabidopsis. Strikingly, plants expressing the phospho-dead version closely resembled *brc1-2* mutants, exhibiting reduced shoot height, enhanced branching, elongated primary roots, and an expanded root apical meristem (RAM), suggesting that loss of phosphorylation renders BRC1 functionally inactive. In contrast, phospho-mimic lines phenocopied BRC1 overexpression, displaying delayed flowering and reduced RAM size, consistent with heightened SL sensitivity (González-Grandío et al., 2017; Wang et al., 2015; Omoarelojie et al., 2019). These findings are consistent with BRC1’s established role in regulating meristem activity via auxin–cytokinin cross-talk (Omoarelojie et al., 2019). Moreover, the delayed flowering phenotype is particularly consistent with BRC1’s known role as a floral repressor through its suppression of FLOWERING LOCUS T (FT) expression (Niwa et al., 2013). Notably, GR24 treatment triggered a dramatic reduction in root growth in phospho-mimic lines, while the phospho-dead variants retained longer roots under identical conditions, reinforcing that the phosphorylation status of BRC1 governs SL responsiveness. This difference likely reflects the reduced RAM size in the phospho-mimic lines, consistent with the expectation that stabilized or overexpressed BRC1 restricts meristem activity. Furthermore, confocal imaging of the lines expressing these GFP-tagged constructs revealed markedly diminished signal in phospho-dead-lines, implying reduced protein stability or enhanced degradation in the absence of phosphorylation. Conversely, the strong but spatially restricted signal in phospho-mimic lines suggests stabilization of the phosphorylated BRC1 form, possibly coupled with localized accumulation in the shortened RAM. These trends align with previous studies implicating BRC1 as a central integrator of SL signals in both shoot and root development (Aguilar-Martínez et al., 2007; Braun et al., 2012).

Together, these findings support a mechanistic model (Figure 8I) in which MPK4-mediated phosphorylation of BRC1 enhances its protein stability and consequently, its action, thereby fortifying SL signaling outputs. The observation that phospho-mimic BRC1 confers hypersensitivity to SL, while phospho-dead BRC1 fails to restore function in the *brc1-2* background, underscores phosphorylation as a critical determinant of BRC1’s biological efficacy within the SL-responsive regulatory framework. Importantly, BRC1, once stabilized, enhances MPK4 expression by directly binding to its promoter, creating a positive feedback loop that amplifies and maintains the SL response. In parallel, MAX2 acts as a molecular executioner that tightly regulates signal spike by targeting unphosphorylated forms of MPK4 and BRC1 for degradation via the ubiquitin–proteasome system. This integrated feedback circuit highlights a previously uncharacterized layer of control in SL signaling and provides new insight into how plants orchestrate hormone-guided developmental processes, particularly shoot branching and overall architecture. Moreover, the above characterized mode of post-translational regulation, involving coordinated phosphorylation and ubiquitination, presents a refined regulatory model (Figure 8I) by which the signaling exerts precise control over key developmental processes such as shoot branching, root architecture, and meristem maintenance.

## Materials and Methods

### Plant Conditions and Treatments

All *A. thaliana* lines used in this study were in the *Columbia-0* (Col-0) background. Mutants *brc1-2* (SALK_091920C), *max2-1* (CS9565), and *mpk4* (CS861181) were obtained from the Arabidopsis Biological Resource Center (ABRC), Ohio State University. Homozygosity of *brc1-2* and *mpk4* was confirmed by PCR genotyping (Supplementary Figure S11 A and B). Seeds of *mpk3*, *mpk6* (Sethi et al 2014, Bhagat et al 2022) and the *proMPK4:GUS* reporter line (Verma et al., 2024) were sourced from laboratory stocks. Seeds were surface sterilized (Lindsey et al., 2017), stratified at 4 °C in the dark for 3 days, and germinated on half-strength Murashige and Skoog (½ MS) medium. Seedlings were grown under short-day conditions (8 h light/16 h dark) at 22 °C with 150 µmol m⁻² s⁻¹ light intensity.

### RNA Extraction

7 day old Arabidopsis seedlings initially grown on half-strength MS agar medium were transferred to liquid half-strength MS medium supplemented with 5 µM GR24 (or DMSO as a control) and incubated at 25 °C with gentle shaking at 50 RPM. Seedlings were treated for 12 hours, and samples were collected at 0, 1, 3, 6, and 12-hour time points by snap-freezing them in liquid nitrogen. Total RNA was isolated from 100 mg of tissue using the Qiagen RNA extraction kit, following the manufacturer’s instructions. To remove any genomic DNA contamination, the extracted RNA was treated with RNase-free DNase I (Qiagen), and the concentration of DNase-treated RNA was quantified using spectrophotometry.

### cDNA Synthesis and Quantitative Real-Time-PCR

First-strand cDNA was synthesized from 2 µg of total RNA using the RevertAid First Strand cDNA Synthesis Kit (Thermo Scientific). The synthesized cDNA was then diluted 20-fold with nuclease-free water, and 1 µL of the diluted product was used in a 10 µL reaction volume for quantitative real-time PCR (qRT-PCR). ACT2 served as the internal reference gene for normalization. Each biological sample was analyzed in triplicate (technical replicates), and the entire experiment was independently repeated three times (biological replicates) to ensure reproducibility.

### Histochemical GUS Staining

For GUS staining, plant tissues were incubated overnight at 37 °C in staining solution containing 50 mM sodium phosphate buffer (pH 7.5), 0.1% (v/v) Triton X-100, 2 mM potassium ferricyanide, 2 mM potassium ferrocyanide, and 1 mg/mL X-Gluc (5-bromo-4-chloro-3-indolyl-β-D-glucuronic acid). To remove chlorophyll, stained plant samples were subjected to sequential ethanol washes with decreasing concentrations (from 90% to 50% v/v), each for 30 minutes. The decolorized samples were then stored in 50% (v/v) ethanol with 10% (v/v) acetic acid until photographed.

### Quantification of GUS Activity

Total protein was extracted from plant samples using GUS extraction buffer containing 10 mM EDTA (pH 8.0), 1% SDS, 50 mM sodium phosphate buffer, 0.1% Triton X-100, 10 mM β-mercaptoethanol, and 1 mM PMSF. Protein concentrations were determined using the Bradford assay (Bradford, 1976). GUS fluorescence was assayed using 5 µg of crude protein extract incubated with 1 mM MUG (4-methylumbelliferyl-β-D-glucuronide). The reaction was terminated at 0, 30 and 60 minutes by adding 0.2 M Na₂CO₃. Fluorescence was measured at an excitation wavelength of 360 nm and emission at 460 nm. GUS activity was expressed as pmol of 4-methylumbelliferone (MU) produced per minute per milligram of protein.

### Immunoblotting Assays

Plant tissues were harvested, flash-frozen, and ground into a fine powder in liquid nitrogen using a pestle and a mortar. The powdered tissue was then resuspended in protein extraction buffer containing 25 mM Tris-HCl (pH 7.5), 15 mM MgCl₂, 15 mM EGTA, 75 mM NaCl, 60 mM β-glycerophosphate, 1 mM DTT, 0.1% NP-40, 0.1 mM Na₃VO₄, 1 mM NaF, 1 mM PMSF and protease inhibitor. The lysate was centrifuged at 13,000g for 10 minutes at 4°C to remove debris, and this step was repeated until the supernatant was clear. Protein was quantified using Bradford assay (Bradford 1976). To denature proteins, samples were mixed with 6× SDS-PAGE loading buffer and boiled for 5-10 minutes. Proteins were separated on SDS-PAGE gels and transferred to nitrocellulose membranes (Bio-Rad) for immunoblotting. Membranes were washed with Tris-buffered saline (TBS) containing 0.1% Tween-20. The following primary antibodies were used at a 1:10,000 dilution: anti-HA (C29F4, CST), anti-AtMPK4 (A6979, Sigma), anti-phospho p42/44 MAPK (Erk 1/2) (Thr202, Tyr204) (9101, CST), anti-AtActin (SAB4301137, Sigma), anti-MYC (C3956, Sigma), anti-phosphothreonine (Santa Cruz, sc-57562), and anti-phosphoserine (Santa Cruz, sc-81514), anti-GST (MA4-004, Invitrogen). HRP-conjugated mouse or rabbit secondary antibodies (Thermo Scientific) were also used at 1:10,000 dilution. Signal detection was performed using Clarity^TM^ Western ECL Substrate (BioRad), and chemiluminescence signals were captured using a iBright 1500 ChemiDoc imaging system.

### Immunoprecipitation Assays

For immunoprecipitation assays, Arabidopsis seedlings expressing *CaMV35S::MPK4-10xMyc, CaMV35S::BRC1-3xHA, CaMV35S::MAX2-10xMyc,* or *CaMV35S::MAX2-3xHA* were harvested, and total protein was extracted as described in the immunoblotting section. Equal amounts of protein extracts were co-incubated in specific combinations (MPK4-Myc with BRC1-HA, BRC1-HA with MAX2-Myc, and MPK4-Myc with MAX2-HA) along with Protein A/G agarose beads (G-Biosciences) for 1 hour at 4°C with gentle rotation to pre-clear nonspecific binding. Approximately 2 mL of each pre-cleared protein mixture was incubated overnight at 4°C with 2 µL of anti-Myc antibody on a rotating wheel. The Myc-tagged proteins (MPK4-Myc or MAX2-Myc) and their interacting partners were then captured using 50 µL of Protein A/G agarose beads, followed by a 4-hour incubation at 4°C with gentle rotation. The beads were collected by centrifugation at 3800 × g for 5 minutes at 4°C and washed three times with 1× PBS buffer to remove unbound proteins. After discarding the supernatant, 2× Laemmli buffer was added directly to the beads. The samples were briefly denatured by heating at 95°C and then subjected to SDS-PAGE. Western blot analysis was performed as described in the previous section.

### *In-silico* Docking Analysis

The predicted three-dimensional structures of BRC1 and MAX2 were obtained from the AlphaFold database, while the structure of MPK4 was sourced from the Protein Data Bank (PDB ID: 7W5C). Protein– protein and protein–DNA docking simulations were performed using the HDock server. Docked models with the highest docking scores were selected for further analysis using the PyMOL standalone software. For the BRC1–MPK4 promoter DNA complex, structural visualization and interaction analysis were conducted using the PDIviz plugin in PyMOL.

### Expression of Recombinant Proteins

For bacterial expression of GST/His-tagged proteins, genes were cloned into pGEX4T2 or pET28a vectors and transformed into *E. coli* BL21. Overnight cultures in 5 mL LB were used to inoculate 300 mL 2×YT, and protein expression was induced with 1 mM IPTG at OD₆₀₀ ≈ 0.5–0.6, followed by incubation at 18°C for 8–12 h. Cells were harvested (5000 rpm, 4°C, 5 min), washed with PBS, and resuspended in lysis buffer (50 mM Tris-HCl pH 7.5, 0.3 M NaCl, 10 mM MgCl₂, 5% glycerol, 1% Triton X-100, 2 mg/mL lysozyme). After 1.5 h on ice, samples were sonicated (6 × 30 s, 70% amplitude) and centrifuged (13,000 rpm, 20 min, 4°C). The supernatant was incubated with GST beads for 4–6 h or overnight at 4°C, followed by 3–4 washes in wash buffer (50 mM Tris-HCl pH 7.5, 0.3 M NaCl, 10 mM MgCl₂, 5% glycerol, 1% Triton X-100). Bound proteins were eluted using wash buffer containing 10 mM reduced glutathione.

### *In vitro* Kinase Assay

*In vitro* phosphorylation of MPK4/MPK4/MPK6-His with BRC1/BRC1^S34A^/ BRC1^S65A^/ BRC1^S34A,S65A^-GST or MBP was carried out following the protocol described by Raghuram et al. 2015. Kinase and substrate were mixed at an enzyme-to-substrate ratio of 1:10 (w/w) and incubated in a reaction buffer containing 20 mM Tris-HCl (pH 7.5), 10 mM MgCl₂, 25 µM ATP, 1 µCi [γ-³²P] ATP, and 1 mM DTT at 28°C for 1 hour. The reaction was terminated by adding 6× SDS loading buffer, followed by boiling the samples for 5 minutes. Proteins were then separated on a SDS-PAGE gel. After electrophoresis, the gel was exposed to a phosphor screen and scanned using a Typhoon FLA 9500 imaging system.

### Yeast One Hybrid Assay

MPK4 promoter fragments were cloned upstream of the Aureobasidin A (AbA) resistance reporter gene in the bait vector pAbAi, generating three constructs: *pAbAi-proMPK4-1, -2*, and -*3*. These constructs were linearized using BstBI and transformed into the Y1H Gold yeast strain. Transformants were selected on SD medium lacking uracil (-Ura), and successful genomic integration of the promoter fragments was confirmed via PCR. The pGADT7-BRC1 plasmid was then introduced into the MPK4 reporter containing yeast strains. Selection was performed on SD medium lacking leucine (-Leu), and BRC1 binding to the MPK4 promoter was assessed by monitoring yeast growth on -Leu medium supplemented with AbA.

### *In vivo* Phosphorylation Assay

*In vivo* phosphorylation of BRC1 was assessed through a mobility shift assay using Phos-tag reagent (NARD Institute), following the protocol outlined by Shi et al. 2022. Protein extracts were prepared from BRC1:HA^Ind^ seedlings treated with either 5 µM GR-24 or 50 µM PD98059, as well as from untreated *brc1-2* and BRC1:HA control lines. For dephosphorylation controls, crude protein extracts were incubated with CIAP for 2 hours. Samples were resolved on 7% SDS-PAGE gels containing 50 µM Phos-tag and 100 mM MnCl₂. Following electrophoresis, gels were washed three times for 10 minutes each in transfer buffer with 10 mM EDTA, followed by a final 10-minute wash in transfer buffer without EDTA. Proteins were then transferred to a nitrocellulose membrane and HA-tagged BRC1 was detected using an anti-HA antibody.

### Yeast Two-Hybrid Assay

Coding sequences (CDS) were cloned into the pGBKT7 and pGADT7 vectors (Clontech) to generate fusion constructs containing the DNA-binding domain (BD) and activation domain (AD), respectively. These constructs were co-transformed in various combinations into the Y2H Gold yeast strain. Positive transformants were initially selected on double dropout (DDO) medium lacking leucine and tryptophan (- Leu, -Trp). For interaction assays, selected colonies were plated onto quadruple dropout (QDO) medium lacking adenine, histidine, leucine, and tryptophan (-Ade, -His, -Leu, -Trp) and incubated at 30°C for 36– 48 hours. Colonies showing robust growth on QDO medium were considered indicative of positive protein– protein interactions. As positive control, we co-transformed yeast with p53-BD and T-antigen-AD (T7-AD), known to interact strongly. For negative controls, Lam-BD (non-interacting bait) and T7-AD (non-interacting prey) were used to rule out non-specific activation.

### ChIP-qPCR

7 day old Col-0 and BRC1-HA^Ind^ seedlings grown on ½ MS agar at 22°C were transferred to ½ MS liquid medium containing 10 µM estradiol (for BRC1-HA^Ind^ induction) or DMSO (Col-0 control) and incubated for 4 hours. Approximately 4 g of tissue per sample was harvested in liquid nitrogen. For chromatin immunoprecipitation (ChIP), samples were crosslinked with 1% (w/v) formaldehyde under vacuum for 15 minutes, then quenched with 2 M glycine for 5 minutes. Tissues were rinsed three times with cold 1× PBS and dried on paper towels before grinding in liquid nitrogen. The powdered tissue was resuspended in nuclei extraction buffer (100 mM MOPS pH 7.6, 10 mM MgCl₂, 0.25 M sucrose, 5% Dextran T-40, 2.5% Ficoll 400, 40 mM β-mercaptoethanol, and protease inhibitors) and incubated for 15 minutes. Nuclei were pelleted by centrifugation at 20,200 × g for 10 minutes at 4°C. Pellets were lysed in nuclei lysis buffer (50 mM Tris-HCl pH 8.0, 10 mM EDTA, 1% SDS, with protease inhibitors) on ice for 30 minutes, followed by dilution in ChIP buffer (initially without Triton X-100). Chromatin was sonicated (5 cycles, 10 s each at ∼P3.2), and then Triton X-100 was added to a final concentration of 1.1%. For immunoprecipitation, 5 µL of ChIP-grade anti-HA antibody was used to pull down HA-tagged chromatin from BRC1-HA^Ind^ and Col-0 samples. DNA-protein complexes were captured using magnetic beads and sequentially washed with low-salt, high-salt, LiCl, and TE buffers. Elution was performed using 0.5× TE buffer, followed by reverse crosslinking with 0.2 M NaCl at 65°C overnight. Samples were treated with proteinase K for 1 hour, and DNA was purified using the Qiagen PCR purification kit. Non-immunoprecipitated chromatin was processed in parallel as input control. Quantitative PCR (qPCR) using promoter-specific primers was conducted, and enrichment was expressed as % Input, calculated using the formula: % Input = 100 × 2^ΔCt^, where ΔCt = [Ct_Input – log₂(input dilution factor)] – CtChIP.

### BiFC and Localisation

For BiFC assays, coding sequences (CDS) were cloned into CD3-1648 and CD3-1651 vectors to generate N-YFP and C-YFP fusion constructs, respectively. For subcellular localization studies, CDS were fused to GFP by cloning into the pGWB5 vector. All constructs were introduced into *Agrobacterium tumefaciens* strain GV3101. Positive clones, along with the p19 suppressor plasmid, were inoculated in YEB medium, followed by incubation at 28°C with shaking. Bacterial cells were harvested, washed with infiltration buffer (10 mM MgCl₂, 10 mM MES, 150 µM acetosyringone), and adjusted to an OD₆₀₀ of 0.5. The cultures were incubated in the dark for 2–3 hours before being infiltrated into the abaxial side of *N. benthamiana* leaves. Leaf discs were collected 2 days post-infiltration and fluorescence was observed using a confocal laser scanning microscope.

### EMSA

DNA probes were generated by PCR amplification using specific primers and subsequently purified with the Qiagen DNA extraction kit. These probes were then radiolabeled and employed in assay to assess binding with bacterially purified BRC1-GST protein, as per the protocol outlined by Singh and Sinha 2016. For the assay, radiolabeled DNA was incubated either alone or with the BRC1-GST protein in a binding buffer composed of 20 mM HEPES (pH 7.4), 10 mM KCl, 0.5 mM EDTA, 0.5 mM DTT, 1 mM MgCl₂, 3% glycerol (v/v), and 1 µg of poly[dI-dC] as a nonspecific competitor. The reaction mixtures were incubated at room temperature for 30 minutes. The resulting DNA-protein complexes were separated on a native 6% polyacrylamide gel using 0.5× TBE as the running buffer and detected via autoradiography using a Typhoon phospho-imager.

### PI Staining

Propidium Iodide (PI) powder (Sigma) was dissolved in PBS to make a 5 mg/mL stock solution, which was then diluted to 5 µg/mL for use. Seedlings were immersed in the working solution for 10 minutes, rinsed with water, and imaged using a Leica TSC SP8 confocal microscope. Imaging was performed using 405 nm excitation and 410 nm emission settings, focusing on Z-sections where quiescent center cells were clearly visible. Root apical meristem length and cell number were quantified using ImageJ software.

### Statistical Analysis

Root and shoot measurements, immunoblot signal intensities, and fluorescence levels were quantified using the ImageJ software (NIH). Statistical analysis was performed using ANOVA or Student’s *t*-test, as specified. Distinct letters denote statistically significant differences at *P*< 0.05. Information on statistical methods and sample sizes for each experiment is provided in the respective figure legends.

### Site Directed Mutagenesis

Site-directed mutagenesis was carried out using Splicing by Overlap Extension PCR (SOEing PCR) as described by Horton et al. (1995).

### Dual Luciferase Assay

MPK4 promoter was cloned into the pGWB635 vector to generate a *pMPK4:luciferase* reporter construct, while BRC1 and its variants were cloned into pGWB5 for GFP-tagged overexpression. All constructs were introduced into *A. tumefaciens* strain GV1301. Positive colonies, along with the p19 helper plasmid, were grown overnight and used for secondary inoculation in YEB medium at 28°C with shaking. Bacterial cultures were pelleted, washed with and resuspended in infiltration buffer (10 mM MgCl₂, 10 mM MES, 150 µM acetosyringone), and adjusted to an OD₆₀₀ of 0.5. After a 2–3 hour incubation in the dark, cultures were infiltrated into the abaxial surface of *N. benthamiana* leaves. Two days post-infiltration, leaves were treated with 5 µL of luciferin (Sigma), and chemiluminescence signals were detected using the iBright1500 ChemiDoc system.

Foe quantification of luciferase activity, leaf tissue was harvested, snap-frozen in liquid nitrogen, and ground to a fine powder 48 hours post-infiltration. Crude protein extraction was performed following the manufacturer’s instructions for the Promega Dual-Luciferase® Reporter Assay System. For each sample, 20 µL of lysate was transferred to a white 96-well plate compatible with luminescence detection. Subsequently, 50 µL of Luciferase Assay Reagent was added to each well, and luminescence was recorded after a 10-second delay using a FLUOstar Omega plate reader to capture the peak signal. Renilla luciferase activity was quantified and used to normalize firefly luciferase activity by calculating the LUC:REN ratio.

### Generation of Double Mutant

The *brc1-2 mpk4* double mutant lines were generated by crossing homozygous *mpk4* and *brc1-2* mutant plants, using *brc1-2* as the female parent. Both the mutants were screened through PCR genotyping to ensure that the mutation was truly homozygous in the plants used for crossing (Figure S11 A and B). A total of 20 F1 progeny with the genotype *brc1-2/BRC1 mpk4/MPK4* were obtained and genotyped using allele-specific primers for both *MPK4* and *BRC1*. These F1 plants were self-fertilized to produce F2 populations in which the mutations segregated. Homozygous *brc1-2 mpk4* double mutants were identified in the F2 generation based on their distinctive phenotype and confirmed by PCR genotyping for both mutant alleles (Figure S11C).

### Production of Stable Transgenic Lines

Thirty-day-old *Arabidopsis thaliana* plants at peak flowering stage, grown under short-day conditions at 22 °C, were used for Agrobacterium-mediated transformation via the floral dip method. Binary vectors containing the gene constructs (MAX2-pGWB14, MAX2-pGWB420, proBRC1-pGWB3, and BRC1^WT/S34A,S65A/S34D,S65D^-pGWB4) were introduced into *A. tumefaciens* strain GV3101. Cultures were grown in YEB medium supplemented with rifampicin and vector-specific antibiotics, harvested at OD₆₀₀ ∼0.6, and resuspended in infiltration medium (5% sucrose, 0.02% Silwet L-77). Plants were inverted and dipped for 1 minute under 10 psi pressure.

Seeds (T0) were surface sterilized (Lindsey et al., 2017) and screened on ½ MS agar containing either 25 mg/L hygromycin or 100 mg/L kanamycin, depending on the vector’s selectable marker. Resistant seedlings were transferred to soil, and T1 seeds were harvested. Segregation analysis on selective media identified transformants with a 3:1 (resistant:sensitive) ratio, indicating heterozygous insertion. T1 lines were advanced, and T2 seeds were screened for homozygosity by selecting lines in which all seedlings grew uniformly under selection, indicating stable, non-segregating transgene insertion. Increased expression of MAX2-HA and MAX2-Myc was confirmed through qRT-PCR (Figure S12 A and B)

## Funding

Authors thank Anusandhan National Research Foundation (File No.: JCB/2020/000041), Department of Science and Technology, Government of India for partial funding.

## Authors’ contribution

LM and AKS planned the study and designed the experiments. LM, NV, DS and SP carried out the experiments. LM and AKS analysed data and wrote the manuscript. All authors read and approved the final manuscript.

## Acknowledgements

LM, NV, and DS are thankful to the Council of Scientific and Industrial Research (CSIR) for the fellowship. NV and SP are thankful to the Department of Biotechnology (DBT) for fellowships. AKS thanks Sir JC Bose Fellowship from the Anusandhan National Research Foundation (File No.: JCB/2020/000041), Department of Science and Technology, Government of India. The authors thank the Radioisotope facility and the Central Instrumentation Facility of NIPGR, New Delhi, India. Authors are thankful to Pratyaksh Dubey for assisting in protein structure prediction using AlphaFold and BRIC-NIPGR for all the support. The authors are thankful to Prof. Pilar Cubas (CNB-CSIC), for providing seeds of Estradiol-inducible *BRC1-HA* and *BRC1-GFP* lines. The authors are also thankful to Prof. Jean Colcombet (INRAE) for providing us with *MPK4-OE* and *MPK4^DE^-OE* lines.

## Declaration of interests

The authors declare no competing interests.

## Declaration of Generative AI and AI-assisted technologies in the writing process

During the preparation of this work the authors used ChatGPT in order to correct grammatical errors. After using this tool/service, the authors reviewed and edited the content as needed and take full responsibility for the content of the publication.

## Supplementary figure legends

**Figure S1.**
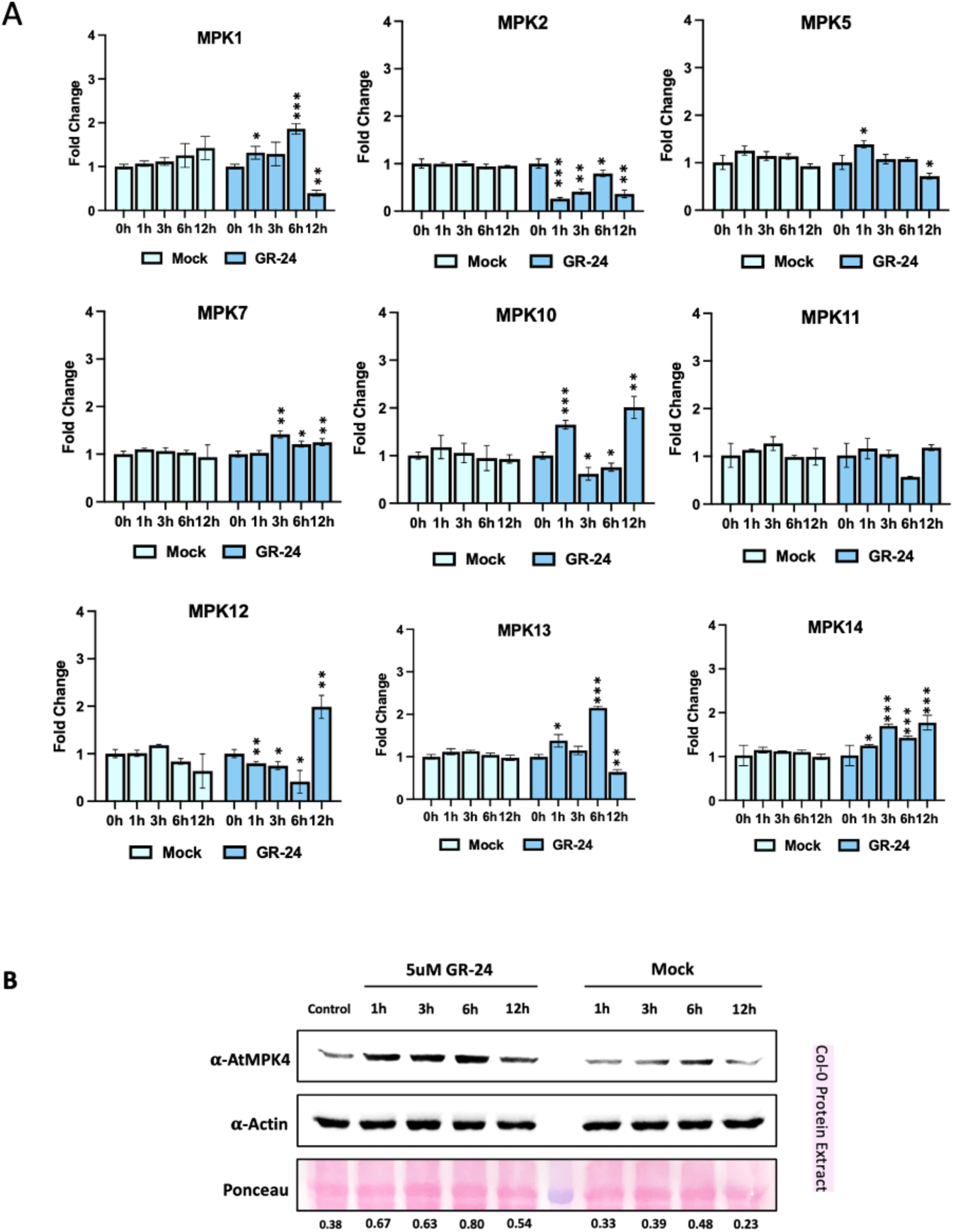
Expression dynamics of MPKs after GR-24 treatment. **A.** Expression analysis of selected group A, B and C MPKs in GR-24 and mock treated, 7day old Col-0 seedlings. Values are means ± s.d. (n= 3). Expression of genes at each time point was compared to 0 h by student’s t test; *p* value < 0.05 (*), < 0.005 (**), <0.0005 (***). **B.** Western blot from crude protein extracts of GR-24 and mock treated, 7day old Col-0 seedlings developed using anti-MPK4 antibody. Ponceau stainied blot shows RuBisCO protein. Actin protein was used as endogenous control. Band intensity was estimated using ImageJ software.

**Figure S2.**
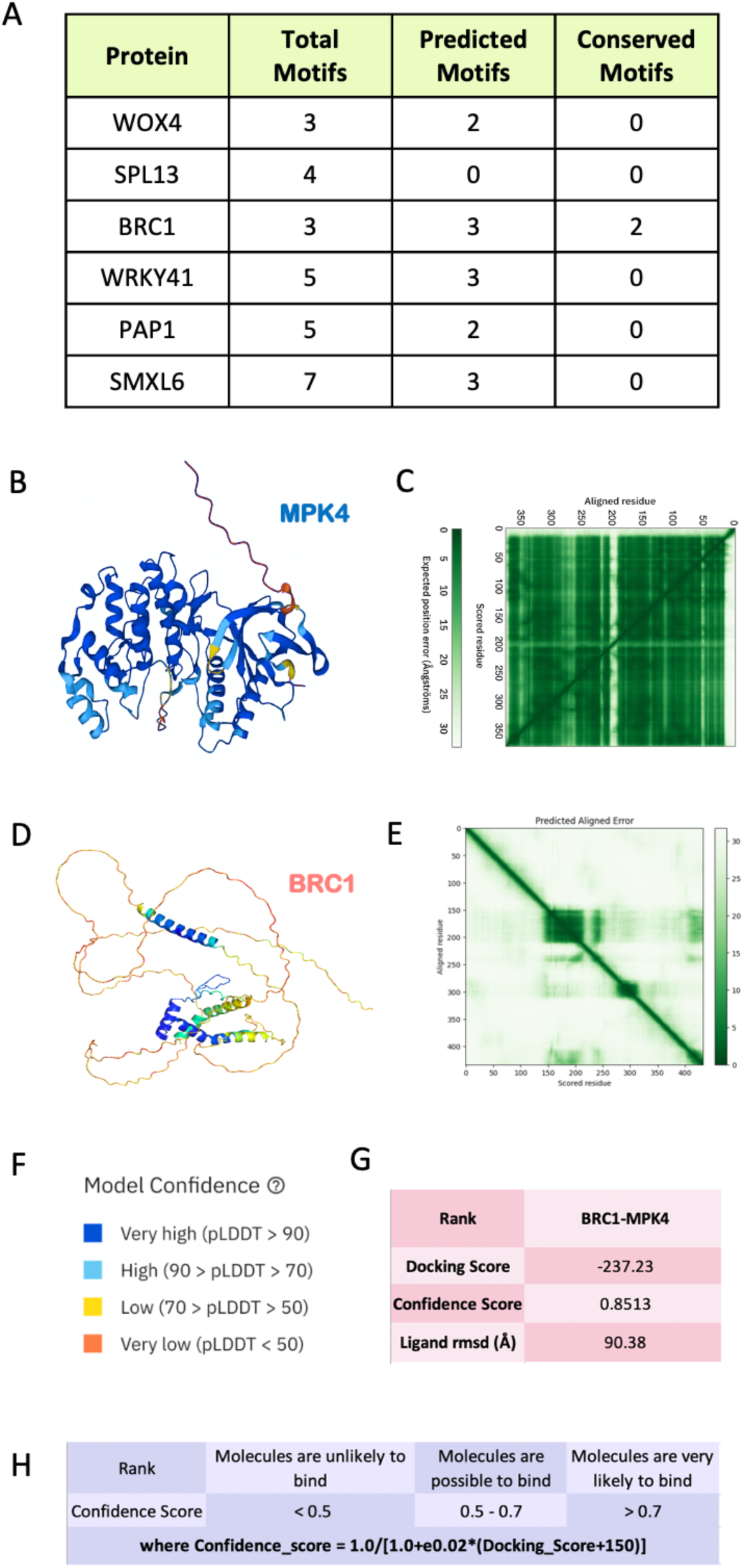
*In silico* analysis of putative MPK4 substrates. **A.** Table showing results of *in-silico* analysis of selected SL response transcription factors for predicting putative phosphorylation SP/TP motifs using NetPhos3.1 tool. Conservation was evaluated by multiple sequence alignment done using Clustal Omega. Protein sequences of genes were retrieved from TAIR. **B.** Predicted 3D structure of MPK4 protein. **C.** Graph showing predicted aligned error of MPK4 protein structure per residue. **D.** Predicted 3D structure of BRC1 protein. **E.** Graph showing predicted aligned error of BRC1 protein structure per residue. **F.** Reference for colour coded pLDDT score annotation. **G.** Table showing confidence score of docked BRC1-MPK4 structure. **H.** Table showing formula for confidence score calculation.

**Figure S3.**
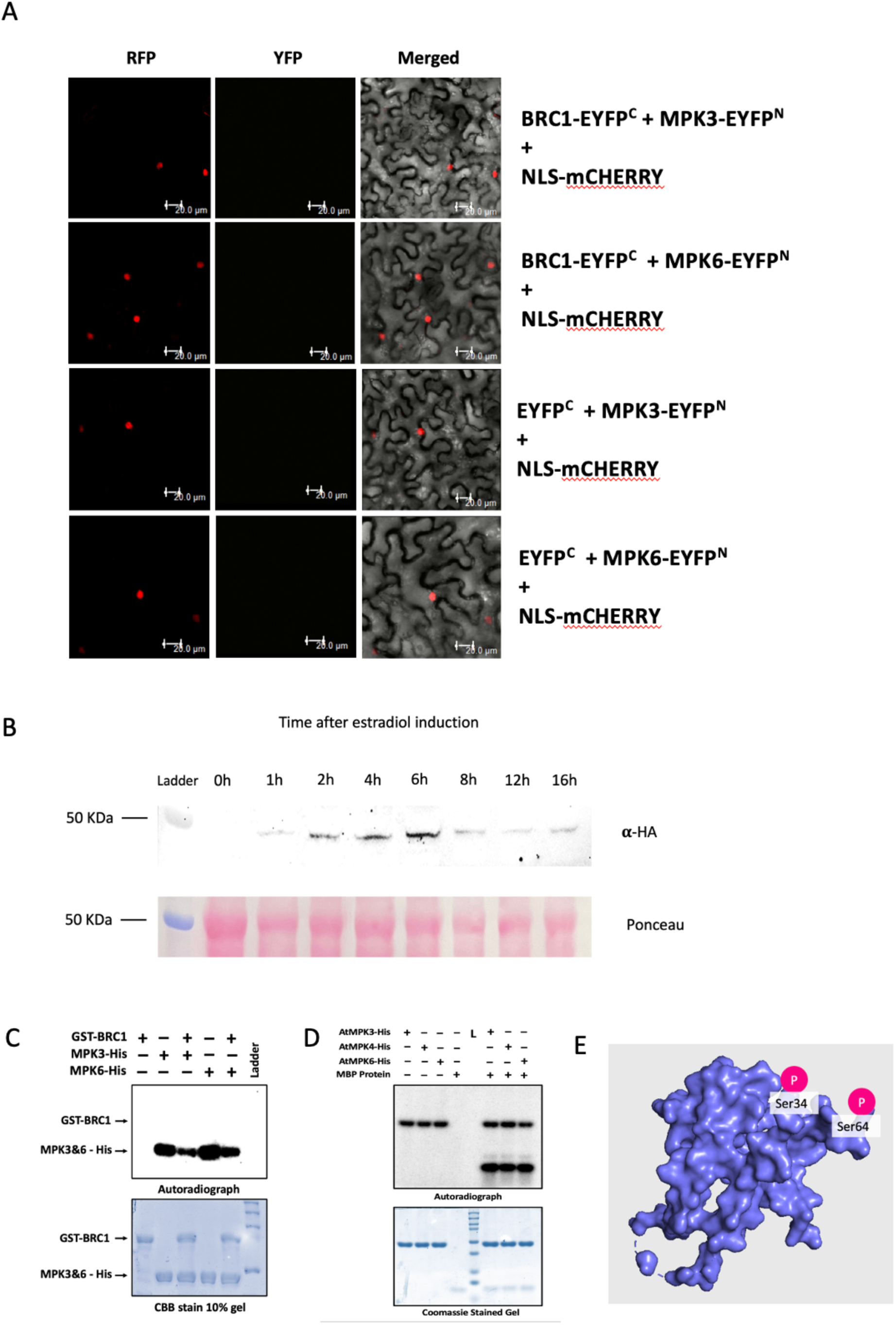
Specificity analysis of MPK4 mediated BRC1 phosphorylation. **A.** Confocal microscopy images of BiFC assay between BRC1 with MPK3 and MPK6 performed transiently in *N.benthamiana* leaves. Scale bar, 20 µm. **B.** Western blot showing time-course analysis of BRC1-HA protein levels in crude protein extract of 7 day old *BRC1*-HA^Ind^ lines post estradiol induction. Blot was developed using anti-HA antibody. Ponceau stained blot shows RuBisCo protein as loading control. **C.** *In-vitro* kinase assay between bacterially-expressed MPK3/MPK6-His and GST-BRC1 proteins. CBB (Commassie Brilliant Blue) stained 10% SDS page gel shows loaded samples **D.** *In-vitro* kinase assay between bacterially-expressed MPK3/MPK6/MPK4-His and MBP proteins showing the activation of MAP kinases. CBB (Commassie Brilliant Blue) stained 10% SDS page gel shows loaded samples **E.** 3D structure of BRC1 protein with positions of MPK4 phosphorylation sites marked using PyMol.

**Figure S4.**
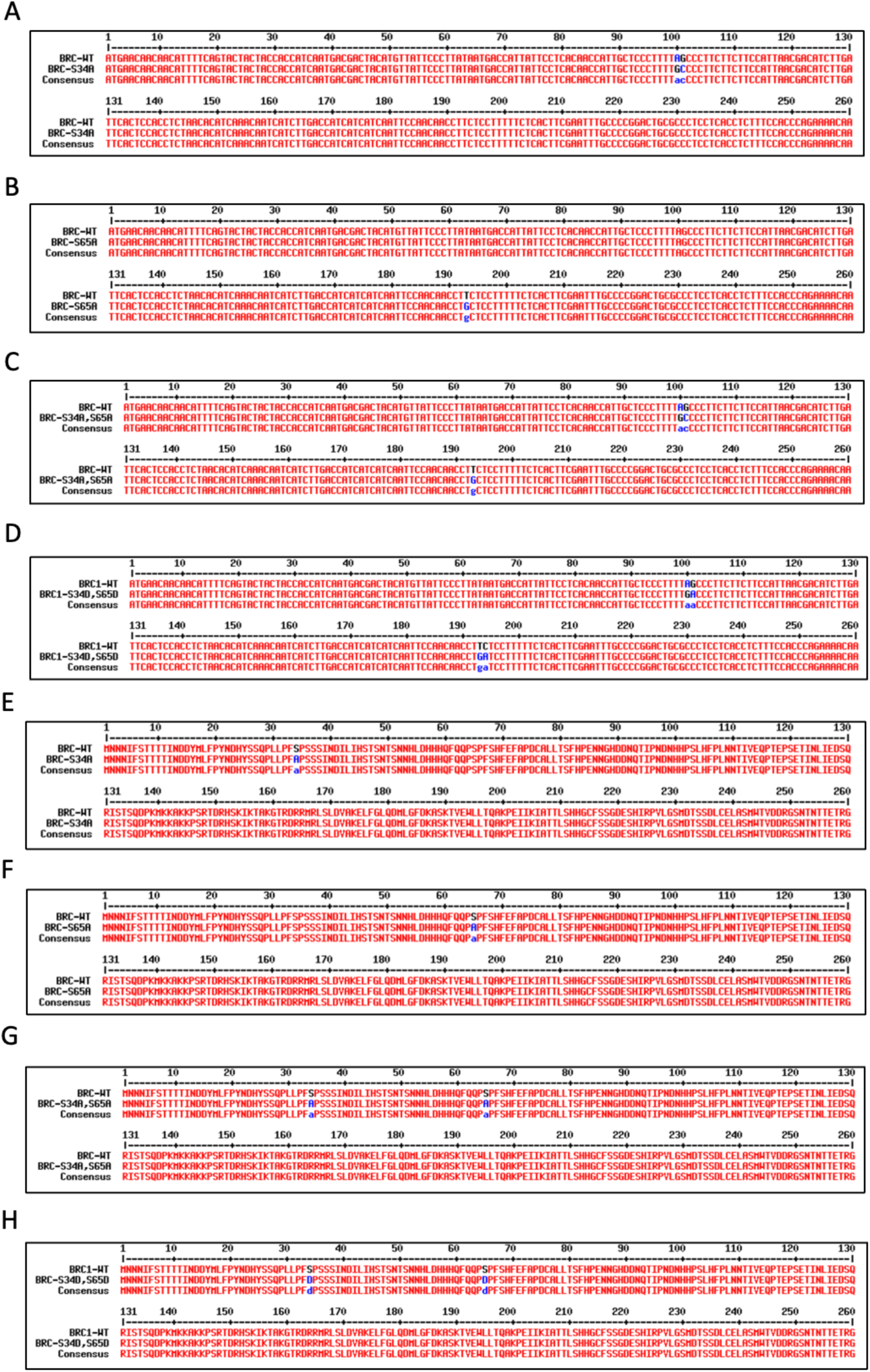
Sequencing results of BRC1 constructs after site-directed mutagenesis. **A.** Snapshot of Multiple sequence alignment (MSA) of BRC1^WT^ and BRC1^S34A^DNA sequences depicting the mis-sense mutation. **B.** Snapshot of MSA of BRC1^WT^ and BRC1^S65A^DNA sequences depicting the mis-sense mutation **C.** Snapshot of MSA of BRC1^WT^ and BRC1^S34A,S65A^ DNA sequences depicting the mis-sense mutations. **D.** Snapshot of MSA of BRC1^WT^ and BRC1^S34D,S65D^ DNA sequences depicting the mis-sense mutations. **E.** Snapshot of Multiple sequence alignment (MSA) of BRC1^WT^ and BRC1^S34A^ protein sequences depicting the mis-sense mutation. **F.** Snapshot of MSA of BRC1^WT^ and BRC1^S65A^ protein sequences depicting the mis-sense mutation **G.** Snapshot of MSA of BRC1^WT^ and BRC1^S34A,S65A^ protein sequences depicting the mis-sense mutations. **H.** Snapshot of MSA of BRC1^WT^ and BRC1^S34D,S65D^ protein sequences depicting the mis-sense mutations.

**Figure S5.**
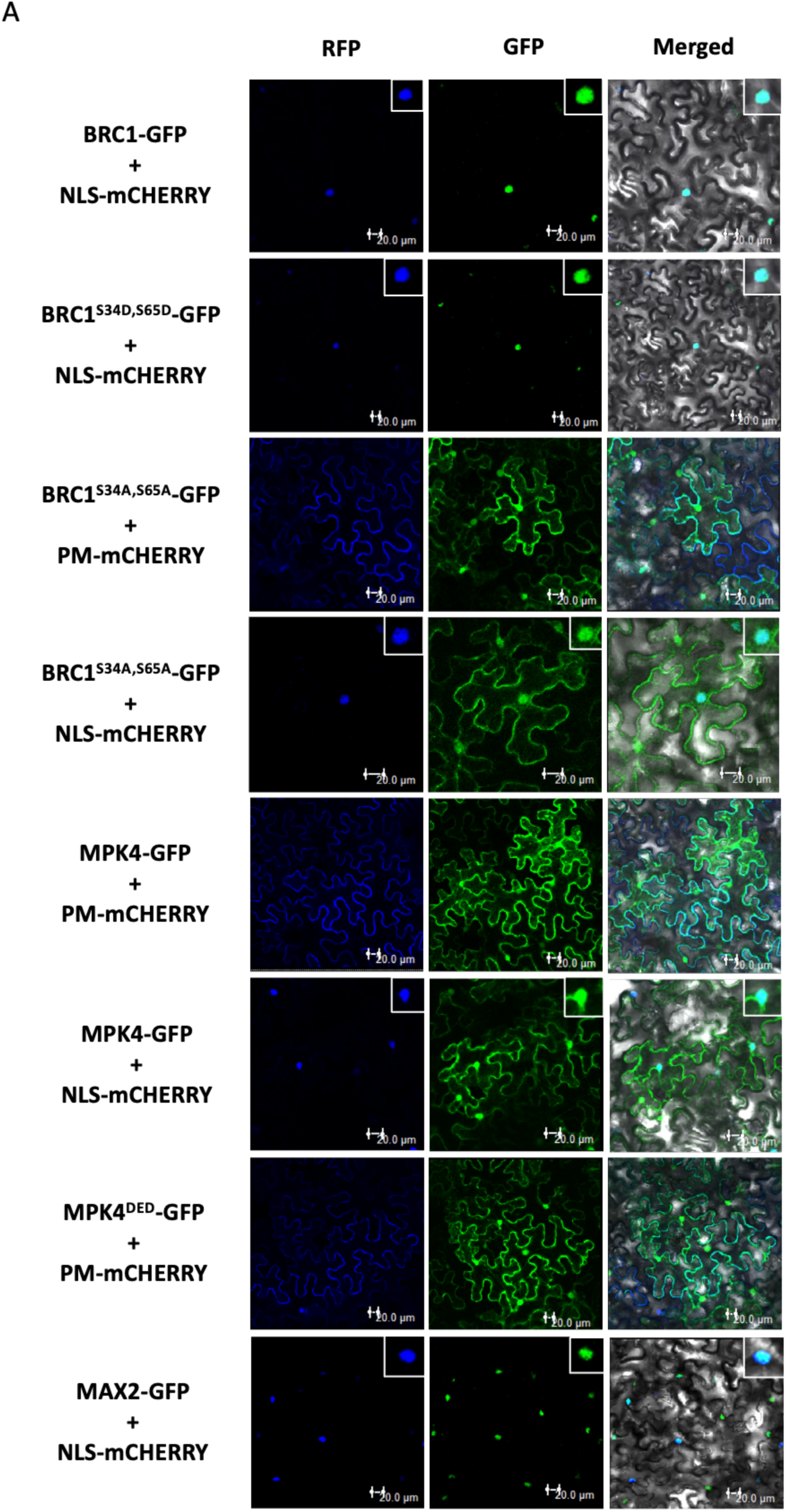
Localisation of BRC1, MPK4 and MAX2 *in planta.* **A.** Confocal microscopy images of transient localization of GFP-tagged proteins in *N.benthamiana* leaves. Inset enlarged image of the signal is present on top right of each image.

**Figure S6.**
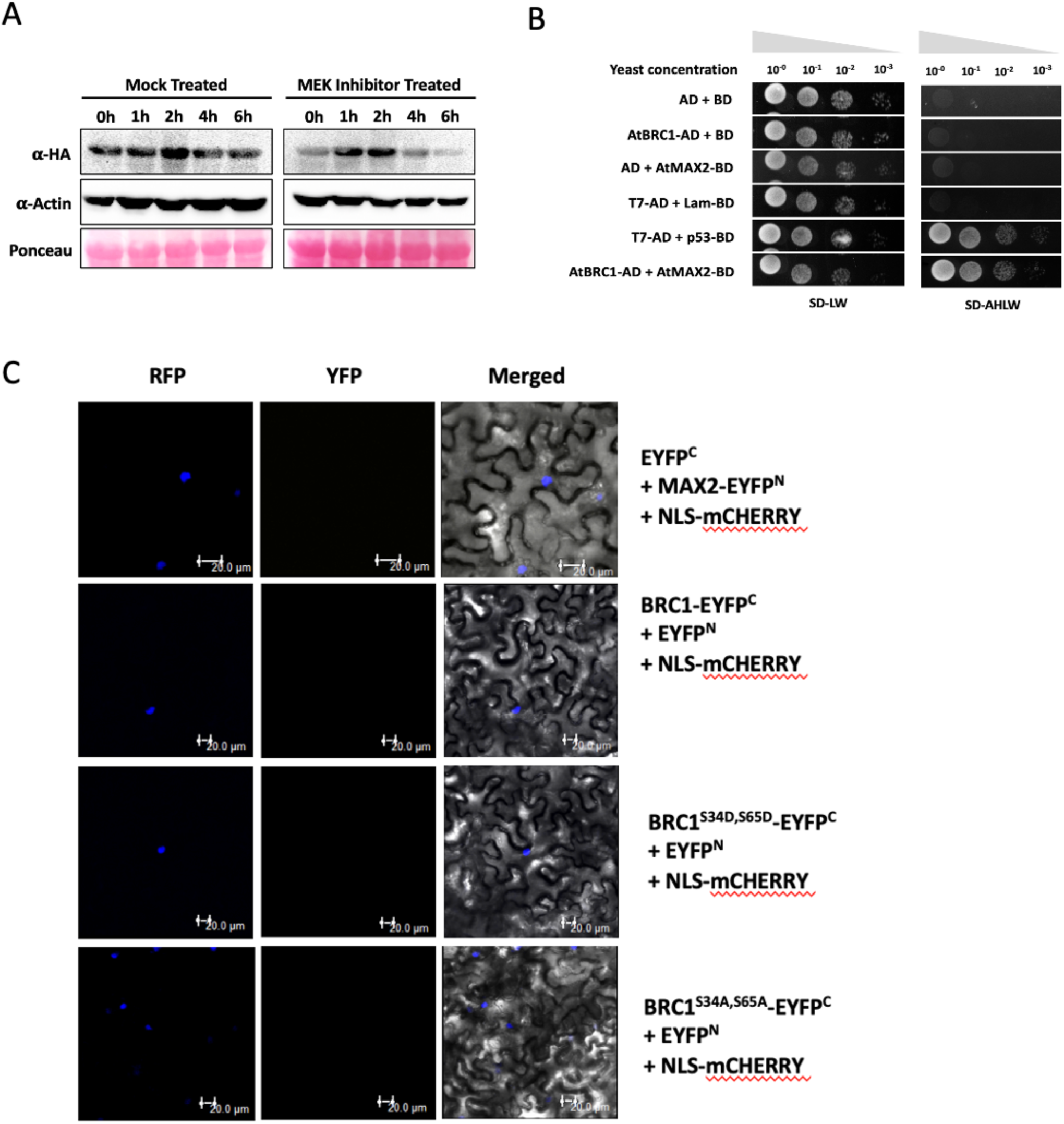
Effect of phosphorylation on stability of BRC1. **A.** Western blot showing time-course analysis of BRC1-HA protein levels in 7day old *BRC1*-HA^Ind^ lines treated with mock/MEK inhibitor. Blot was developed by anti-HA antibody. Ponceau stained blot shows RuBisCo protein as loading control. **B.** Yeast-two-hybrid assay of MAX2 with BRC1. SD-LW and SD-AHLW stand for supplement dropouts -leucine, -tryptophan and -adenine,-histidine,-leucine-tryptophan respectively. **C.** Confocal microscopy images of negative controls used in BiFC assay between BRC1 and MAX2 performed transiently in *N.benthamiana* leaves. Scale bar, 20 µm.

**Figure S7.**
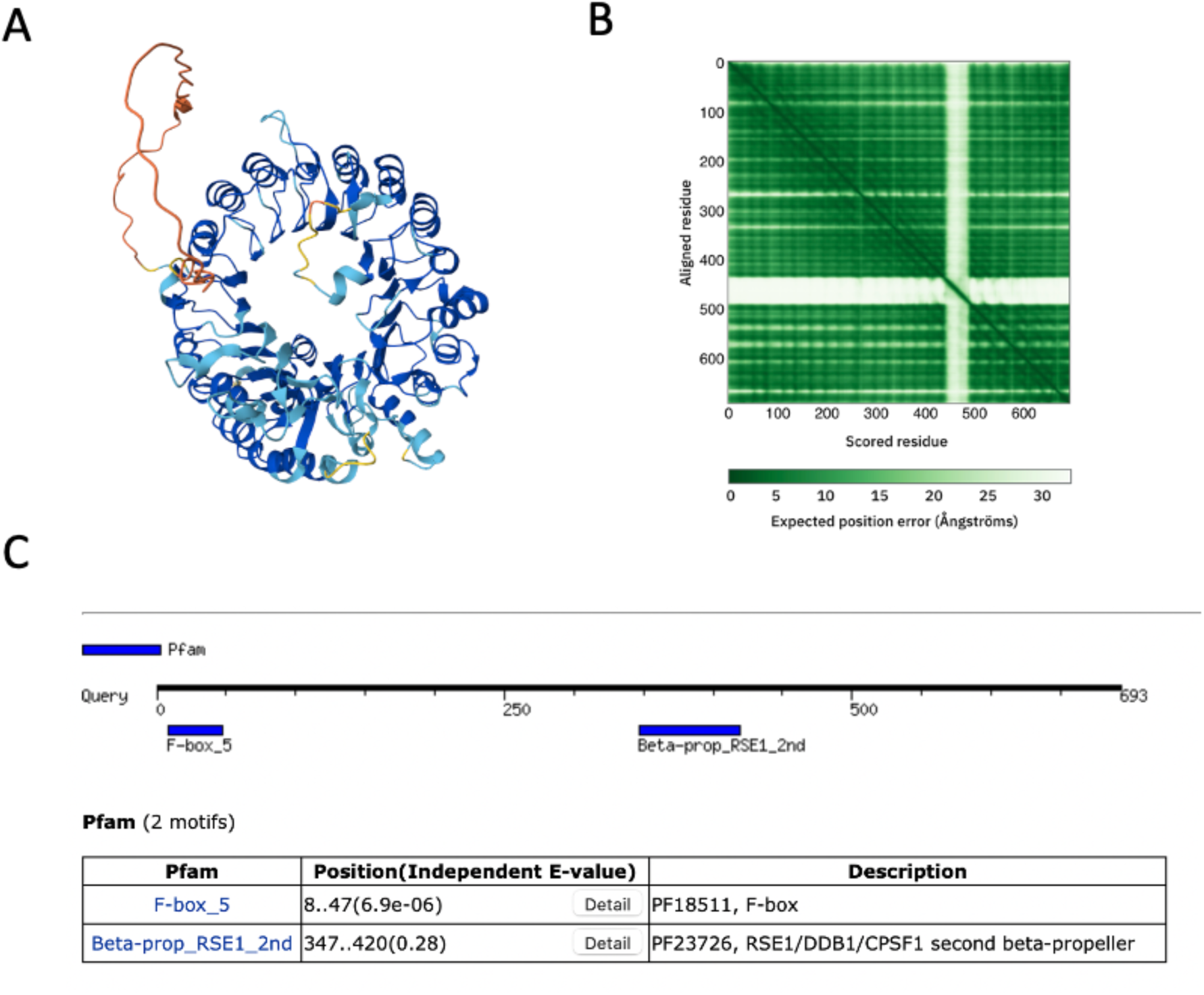
*In silico* analysis of MAX2 protein. **A.** Predicted 3D protein structure of MAX2. **B.** Graph showing predicted aligned error of the MAX2 protein structure per residue. **C.** Predicted motifs in MAX2 protein sequence using Motif Finder tool.

**Figure S8.**
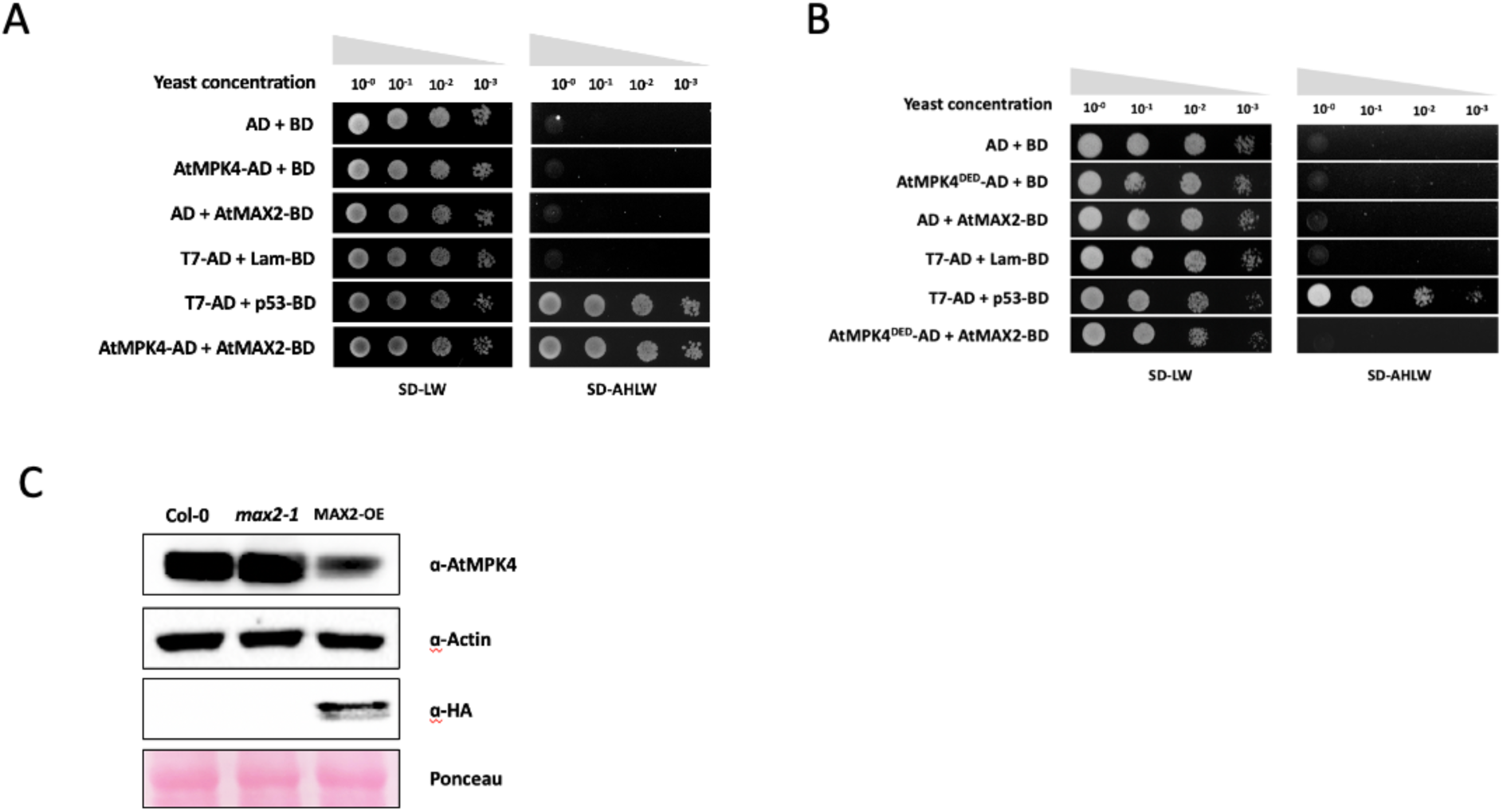
Effect of MAX2 on MPK4 protein. **A.** Yeast-two-hybrid assay showing interaction of MAX2 with MPK4. SD-LW and SD-AHLW stand for supplement dropouts -leucine, -tryptophan and -adenine, -histidine, -leucine, -tryptophan, respectively. **B.** Yeast-two-hybrid assay of MAX2 with MPK4DED. SD-LW and SD-AHLW stand for supplement dropouts -leucine, -tryptophan and -adenine,-histidine,- leucine - tryptophan respectively. **C.** Western Blot using protein crude extracts of 7day old Col-0, *max2-1* and *MAX2*- OE (*MAX2*-HA) lines. Blots were developed using anti-MPK4 antibody. Actin was used as loading control. Blot developed with anti-HA antibody depicts protein levels of MAX2 in *CaMV35S::MAX2:*3xHA line.

**Figure S9.**
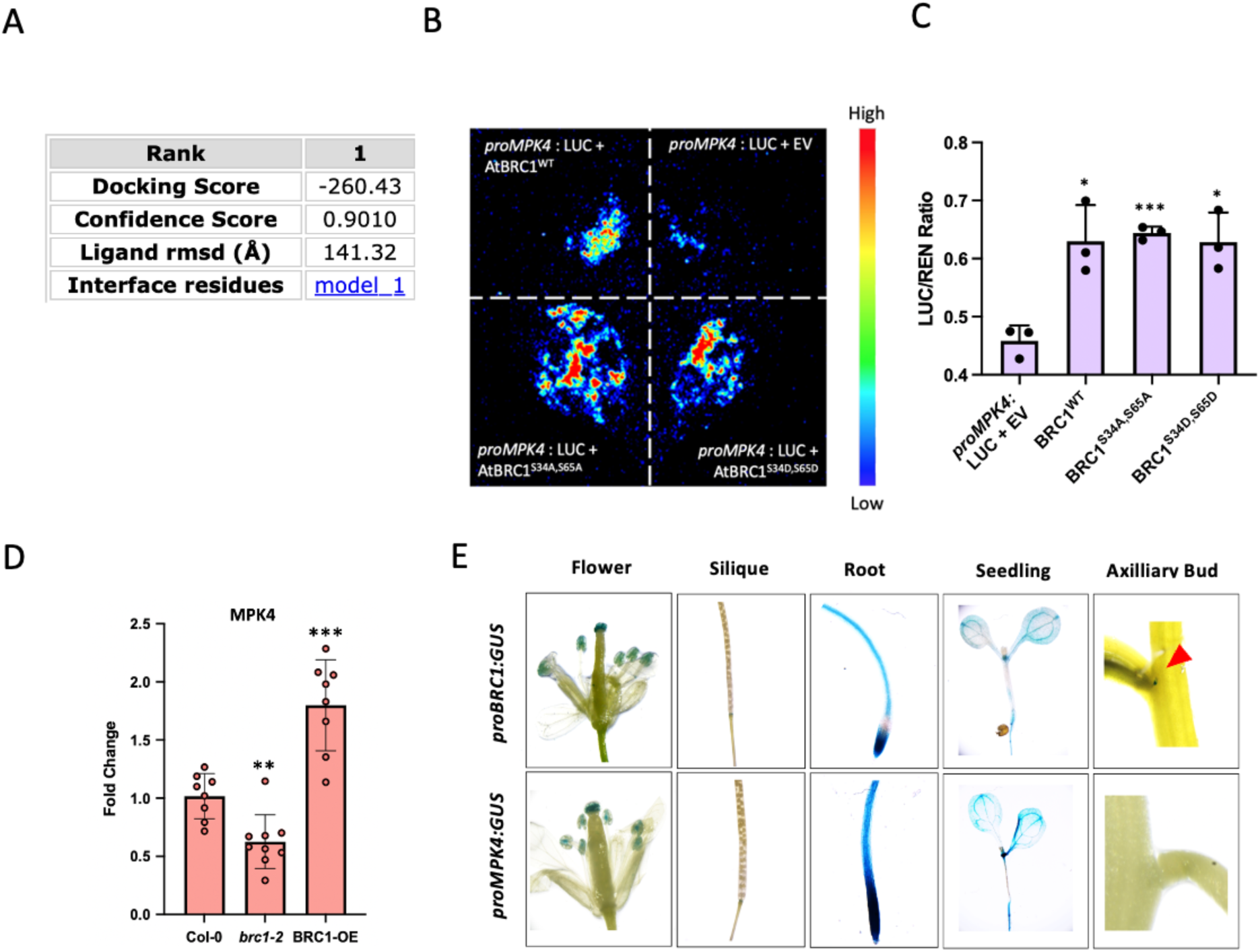
Interaction of BRC1 with MPK4 promoter. **A.** Table showing confidence score of docked structure of BRC1 and *MPK4* promoter. **B.** Luciferace assay showing interaction between BRC1 variants and *pMPK4-2* of transiently infiltrated in *N. benthamiana* leaf. **C.** Quantification of Renilla luciferase activity in protein crude extracts of transiently infiltrated *N. benthamiana* leaves from experiment in figure S12B using luciferin as substrate. Values are means ± s.d. (n= 3). Significant difference was calculated by Student’s t test; *p* value < 0.0005 (***). **D.** Transcript level of *MPK4* in 7day old Col-0, *brc1-2* and *BRC1*-OE (*BRC1*-HA^Ind^) seedlings. Values are means ± s.d. (n= 9). Expression of MPK4 in *brc1-2* and *BRC1*- HA^Ind^ seedlings was compared to Col-0 using Student’s t test: *p* value < 0.005 (**), <0.0005 (***).. **E.** Histochemical GUS staining showing activities of *MPK4* and *BRC1* promoters in different parts of *proMPK4:GUS* and *pro:BRC1:GUS* transgenic lines, respectively.

**Figure S10.**
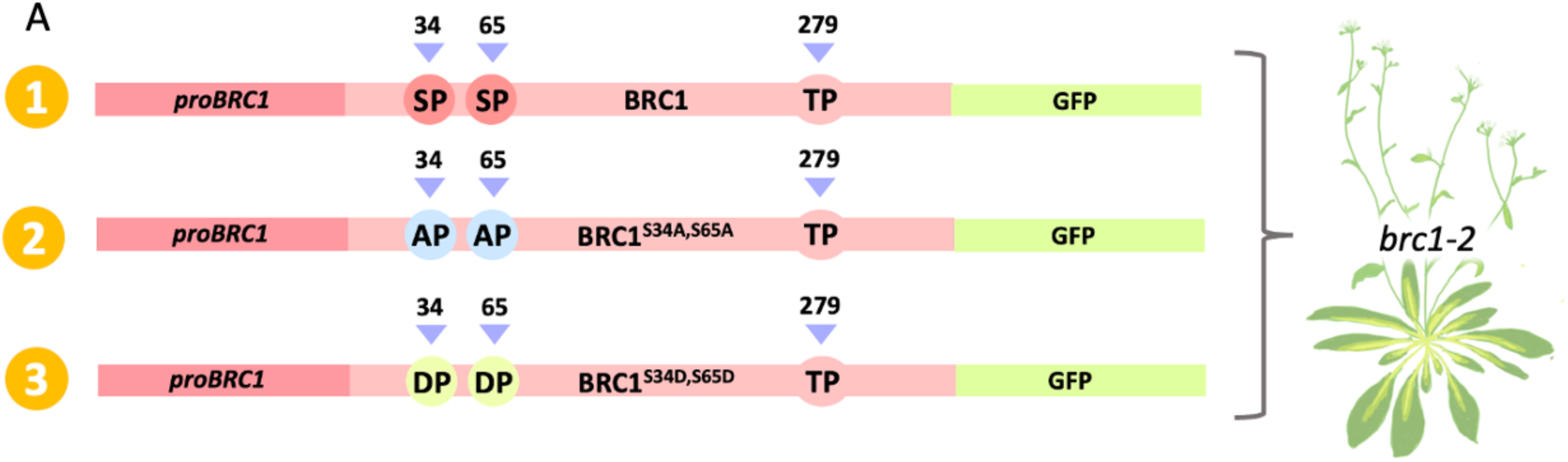
Image depicting constructs used for creating BRC1 complementation lines. **A.** Pictorial representation of constructs having WT, phosphodead (S34A,S65A), and phosphomimic (S34D,S65D) variants of *BRC1* under native promoter. These constructs were used to complement *brc1-2* mutant to generate WT, phosphodead and phosphomimic *BRC1* complementation lines.

**Figure S11.**
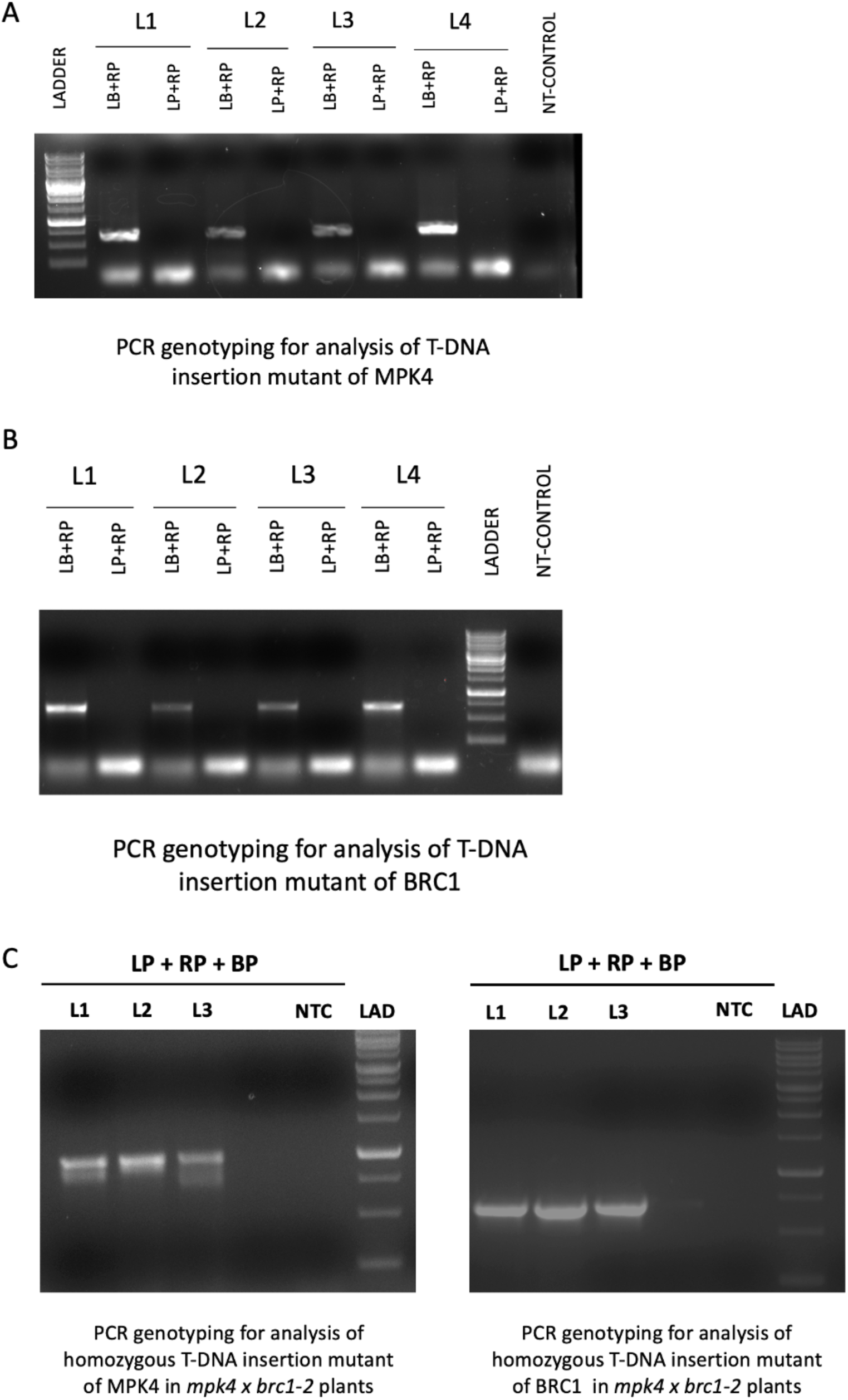
PCR genotyping of *brc1-2, mpk3* and double mutant. **A.** PCR-genotyping for screening of T-DNA insertion mutant of *MPK4*. 1Kb ladder was used to estimate band size. NT stands for no template control. **B.** PCR-genotyping for screening of T-DNA insertion mutant of *BRC1*. 1Kb ladder was used to estimate band size. NT stands for no template control. **C.** PCR-genotyping for screening of T-DNA insertion mutant of *MPK4* and *BRC1* in *mpk4 x brc1-2* double mutant generated by a cross between *mpk4* and *brc1-2* mutants. 1Kb ladder was used to estimate band size. NTC stands for no template control.

**Figure S12.**
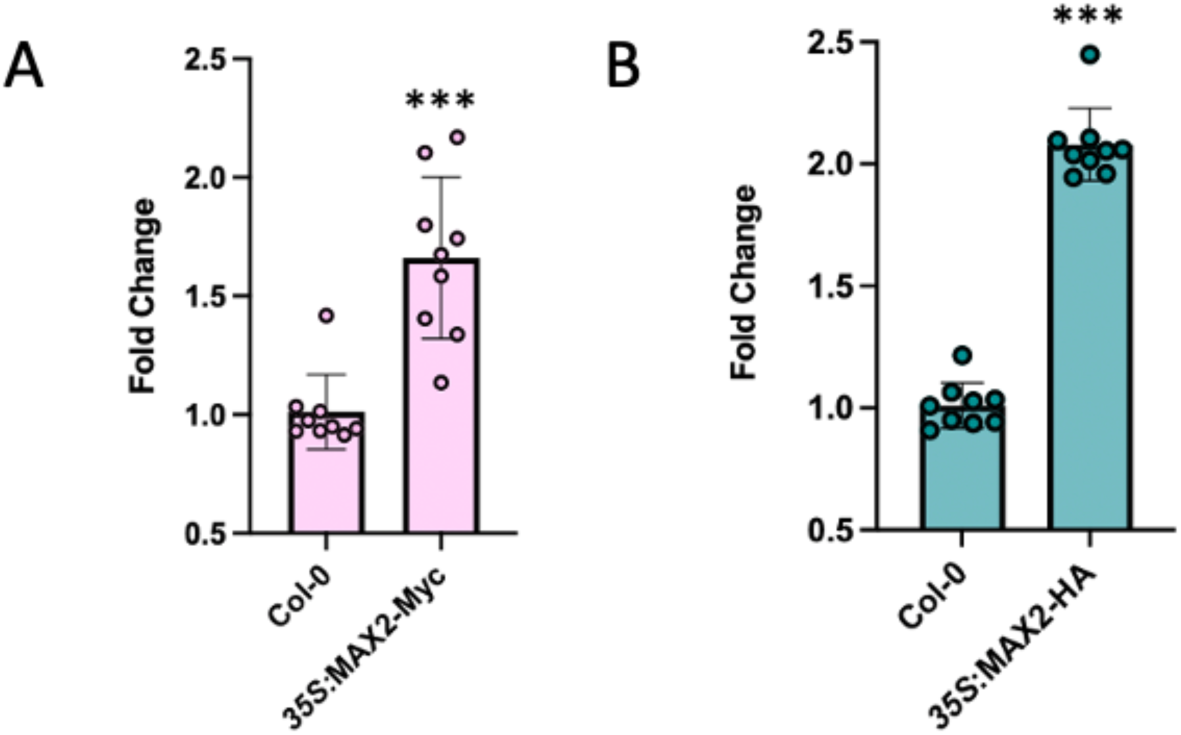
Expression analysis of MAX2 in its overexpression lines. **A.** Real-Time PCR of *MAX2* in *35S:MAX2-Myc* overexpression plants generated using Agrobacterium mediated transformation. RNA was extracted from 7day old seedlings. Values are means ± s.d. (n= 9). Expression of *MAX2* in *35S:MAX2-Myc* plants was compared to Col-0 using Student’s t test: *p* value < 0.0005 (***). **B.** Real-Time PCR of MAX2 in *35S:MAX2-HA* overexpression plants generated using Agrobacterium mediated transformation. RNA was extracted from 7day old seedlings. Values are means ± s.d. (n= 9). Expression of *MAX2* in *35S:MAX2-HA* plants was compared to Col-0 by Student’s t test: *p* value < 0.0005.

## Notes

### Competing Interest Statement

The authors have declared no competing interest.

